# Global loss of responsiveness in key regulator metabolites and elevated enzyme proteins as metabolic dysregulation in skeletal muscle and liver of obese mice during starvation

**DOI:** 10.1101/2025.01.22.634184

**Authors:** Dongzi Li, Keigo Morita, Toshiya Kokaji, Atsushi Hatano, Akiyoshi Hirayama, Tomoyoshi Soga, Yutaka Suzuki, Masaki Matsumoto, Takaho Tsuchiya, Haruka Ozaki, Satoshi Ohno, Hiroshi Inoue, Yuka Inaba, Hikaru Sugimoto, Yifei Pan, Shinya Kuroda

## Abstract

Starvation induces complex metabolic adaptations in skeletal muscle, a key tissue for maintaining energy homeostasis; however, these adaptations are largely impaired in obesity. How obesity alters global metabolic adaptations to starvation in skeletal muscle remains unclear. Here, we analyzed the metabolic adaptations on a trans-omics scale during starvation in skeletal muscle from wild-type (WT) and leptin-deficient obese (*ob*/*ob*) mice. We measured multi-omics data during starvation and constructed global trans-omics networks in WT and *ob*/*ob* mice. We found that starvation induces “responsiveness” in WT mice, characterized by increases or decreases in key regulator metabolites, including ATP and AMP, as well as enzyme proteins, leading to global regulation of metabolic pathways, which was lost in *ob*/*ob* mice. In contrast, during starvation, *ob*/*ob* mice exhibit “difference” in comparison to WT mice, manifested by the persistently elevated expression of metabolic enzymes. These features were similarly found in liver, another key metabolic organ. Thus, global loss of responsiveness and elevated enzyme proteins are systemic features of metabolic dysregulation in *ob*/*ob* mice.

## Introduction

Starvation, a state of prolonged nutrient deprivation, triggers a series of metabolic adaptations that are critical for survival.^1^ During this period, the body must carefully manage energy reserves to maintain essential physiological functions.^2^ The adaptive response to starvation, as well as the therapeutic potential of starvation has been widely studied across various organisms, including yeast^3^, *C. elegans*^4-7^, fish^8-10^, mice^11-15^, rats^16-18^, and humans^19-21^, providing insights into conserved mechanisms of metabolic homeostasis. Among mammals, skeletal muscle, liver, and adipose tissue are three key energy-related tissues that play central roles in the adaptation to starvation.^22-26^ Skeletal muscle serves as a reservoir of amino acids^23^, mobilized for gluconeogenesis in liver to maintain blood glucose levels^27^, supporting energy needs in critical organs like the brain.^28^ Previous study has examined time-course multi-omics data in the liver during starvation, revealing global patterns of metabolic adaptations over time^29^, yet similar analyses in skeletal muscle remain unavailable.

Metabolic adaptation during starvation is characterized by changes in molecular concentrations over time, which is termed “responsiveness” in this study. In skeletal muscle, starvation triggers a decrease in glycogen content^30-34^, an increase of gene expression related to lipid metabolism^35-37^, and a decrease in protein synthesis^38, 39^. Studies in the liver have revealed that starvation decreased glycogen stores^30, 33, 34^, increased gluconeogenic gene expression^40-42^, and increased both fatty acid oxidation and ketogenesis^43-45^. While these studies reveal key molecular responsiveness to starvation in skeletal muscle and liver, the limited number of time points analyzed has restricted our ability to capture a dynamic, continuous picture of the adaptation process to starvation.

Metabolic inflexibility associated with obesity is characterized by “differences” in molecular concentrations between healthy and obese individuals. Human study found that obese individuals had a lower capacity for fat oxidation and displayed inflexibility in regulating fat oxidation during starvation in skeletal muscle.^46^ Another study reported that the shift from glucose to lipid oxidation was blunted in obese subjects.^47^ In obese rats, abnormally higher activation of the mammalian target of rapamycin (mTOR) pathway was observed in the liver and skeletal muscle during starvation, compared with lean controls^48^. These findings highlight the impaired adaptive mechanisms in specific pathways in obese subjects, suggesting metabolic dysregulations during starvation compared to lean counterparts; however, these individual observations fall short of providing a comprehensive view of how obesity affects the overall metabolic adaptations to starvation.

To overcome these limitations, trans-omics analysis—an integrative approach combining metabolomics, transcriptomics, and proteomics—provides a comprehensive approach to uncovering the regulatory networks underlying metabolic adaptations to starvation. Through this approach, we can analyze how tissues adapt to starvation by examining the interplay between different molecular layers. In metabolic regulation, metabolites serve not only as substrates and products in reactions but also as cofactors and allosteric regulators of the enzymes that catalyze these processes.^49-51^ The activity of these enzymes is further regulated by their levels—often dictated by gene expression changes—and by post-translational modifications, such as phosphorylation, which alter their functional state.^52, 53^ These regulatory layers are coordinated by transcription factors and signaling molecules, which, through their own activation and modifications, influence both enzyme activity and gene expression. Consequently, metabolic reactions are governed by an intricate network, integrating metabolites, enzymes, transcription factors, and signaling molecules, all working together to enable dynamic metabolic adaptation. Previous studies in our lab have applied trans-omics analysis to the liver during starvation^29^, revealing that the loss of responsive key regulator metabolites such at ATP and AMP caused the temporal dysregulation in *ob*/*ob* mice.

Among the various metabolites involved in metabolic regulation, energy-related metabolites play a particularly crucial role during shifts in energy status. These metabolites, including ATP, ADP, AMP, NAD, and SAM, participate in energy transfer, redox reactions, and regulatory processes. Previous studies from our lab have highlighted their significance in energy-related adaptations. During starvation, a state of energy deficiency, liver samples demonstrated responsiveness among energy-related metabolites.^29^ Similarly, during oral glucose tolerance tests (OGTT), which simulate energy intake, both liver and muscle samples exhibited responsiveness among these metabolites.^54, 55^ Notably, the AMP/ATP ratio, a key indicator of cellular energy status^56^, is detected by the AMP-activated protein kinase (AMPK) signaling pathway, a well-established mechanism for coordinating energy balance. ^57-59^ Activation of AMPK has been shown to promote energy-generating processes, such as fatty acid oxidation and glucose uptake, while inhibiting energy-consuming processes, including protein synthesis and lipid biosynthesis, thereby helping cells adapt to energy scarcity during starvation.^60^

In this study, we analyzed metabolomic, transcriptomic, and proteomic data, as well as Western blot of protein phosphorylation, from skeletal muscle during a 24-hour starvation period, utilizing trans-omics networks to highlight responsiveness and differences between WT and *ob*/*ob* mice, a widely used model for investigating obesity and its metabolic consequences. This methodology enabled the identification of responsiveness to starvation and differences between normal and obese skeletal muscle during starvation. We observed a global loss of responsiveness and elevation of enzyme proteins in the skeletal muscle of *ob*/*ob* mice. Additionally, we compared metabolic dysregulation associated with obesity in skeletal muscle and liver, revealing the similar patterns of disruption in skeletal muscle and liver of *ob*/*ob* mice. Our findings demonstrate that the global loss of responsiveness and the elevation of enzyme proteins in key metabolic organs, such as skeletal muscle and liver, are systemic features of obesity-related metabolic dysregulation, offering valuable insights for developing therapeutic strategies for obesity-associated metabolic disorders.

## Results

### Workflow overview: Trans-omics analysis of metabolism during starvation in skeletal muscle of WT and *ob*/*ob* mice and its comparison with liver

To uncover the global adaptation patterns to starvation in healthy skeletal muscle and the pathological changes associated with obesity, we constructed and analyzed trans-omics networks in WT and *ob*/*ob* mice during starvation (Fig. 1). We collected multi-omics data from mice experiments, measuring the metabolome, transcriptome, and proteome in skeletal muscle during starvation (Fig. 1a). Additionally, we measured several phosphoproteins by western blot. To assess the effectiveness of starvation, we confirmed that blood insulin decreased in both WT and *ob*/*ob* mice while blood glucose decreased in WT mice (p < 0.05) (fig. S1, Data File S1).

**Fig. 1:**
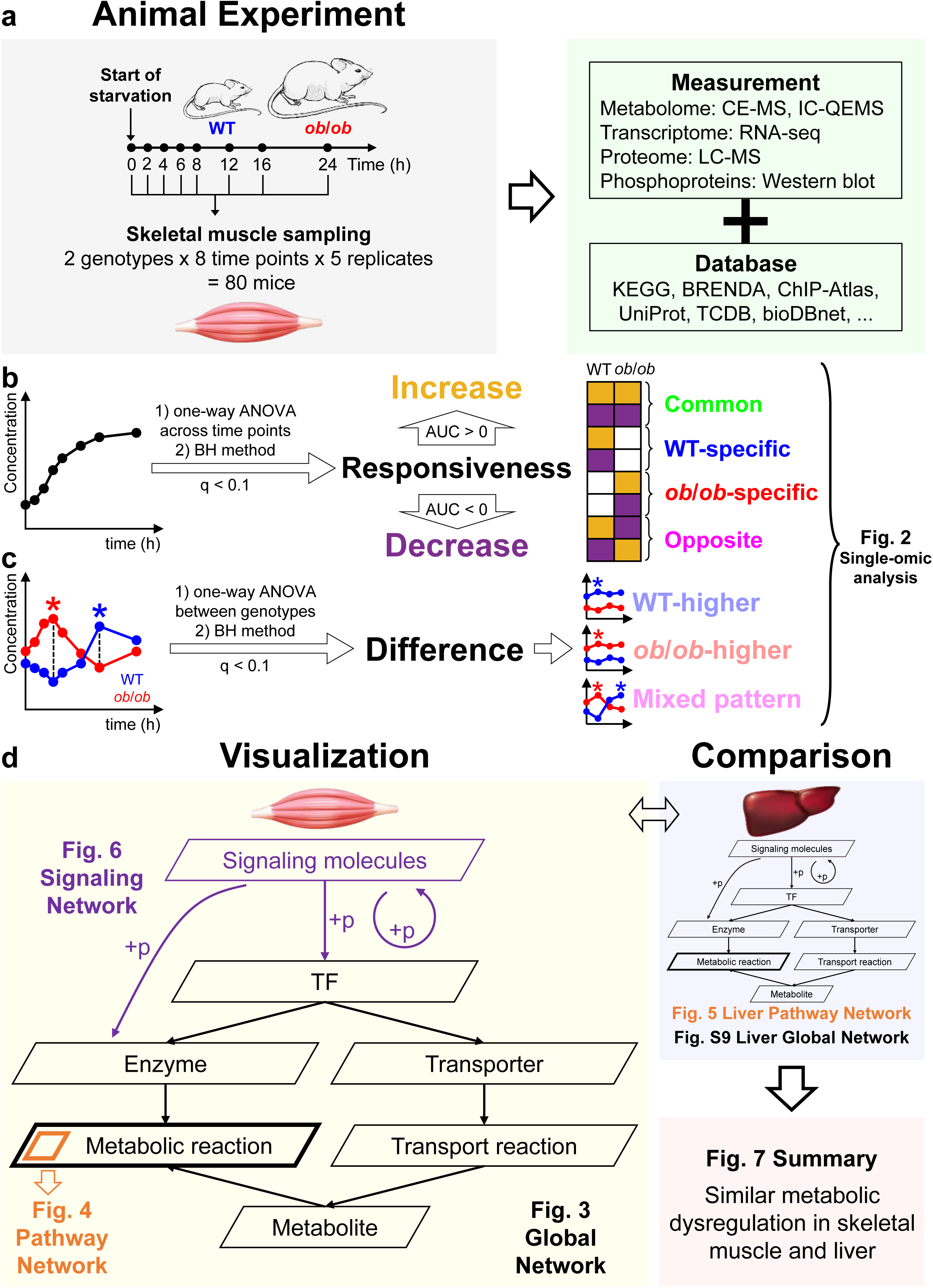
Overview of the study. (a) Workflow of the study, from animal experiments to omics data analysis. At each time point, gastrocnemius muscle samples were collected from five WT and five *ob*/*ob* male mice. (b) Schematic illustrating the definition of responsiveness. Responsiveness was defined as q < 0.1 across time points, determined using one-way ANOVA with BH correction. (c) Schematic illustrating the definition of difference. Difference was defined as q < 0.1 at any time point between WT and *ob*/*ob* mice. (d) Visualization of global, pathway, and signaling networks in skeletal muscle, followed by comparative analysis with liver networks to identify systemic features.

From the measured data, we performed a single-omic layer analysis to identify molecular features during starvation. We categorized these features into two types: “responsiveness” and “difference”. “Responsiveness” was defined as changes across time points. Starvation-responsive molecules were identified by one-way ANOVA (ANOVA-like testing of edgeR^61-63^ for the transcriptome) to compare different time points. The Benjamini-Hochberg (BH) method was used for false discovery rate (FDR) adjustment. Molecules with q values less than 0.1 were defined as starvation-responsive. Based on the area under the curve (AUC) of the log2-normalized time courses, molecules with positive AUCs were classified as increased, while those with negative AUCs were classified as decreased. Starvation-responsive molecules were classified into four groups: “common” (responsive in both WT and *ob*/*ob* mice), “WT-specific” (responsive only in WT mice), “*ob*/*ob*-specific” (responsive only in *ob*/*ob* mice), and “opposite” (showing opposite responses between WT and *ob*/*ob* mice). “Difference” was defined as differential levels between WT and *ob*/*ob* mice during starvation. Starvation-differential molecules were identified by one-way ANOVA (ANOVA-like testing of edgeR for the transcriptome) to compare WT and *ob*/*ob* mice at each time point. The BH method is used for FDR adjustment. Molecules with q values less than 0.1 at any time point were defined as starvation-differential. Starvation-differential molecules were classified into three groups: “WT-higher” (where all differential time points showed higher levels in WT mice), “*ob*/*ob*-higher” (where all differential time points showed higher levels in *ob*/*ob* mice), and “mixed pattern” (higher in WT mice at some time points but higher in *ob*/*ob* mice at others).

Using these starvation-responsive and -differential molecules, we constructed starvation-responsive and -differential trans-omics networks, respectively (Fig. 1d). The networks include three types: global networks, pathway networks and signaling networks. We also re-analyzed the pathway networks in the liver based on our previous study of starvation.^29^ Through these analyses, we elucidated metabolic dysregulations during starvation associated with obesity in both skeletal muscle and liver, two major metabolic organs.

### Identification of starvation-responsive and -differential molecules by single-omic analysis in the skeletal muscle of *ob*/*ob* mice

We measured three omics layers—metabolome, transcriptome, and proteome—in skeletal muscle and performed single-omic layer analysis in WT and *ob*/*ob* mice (Fig. 2). Molecules detected in less than half of the replicates (fewer than 3 out of 5) at any time point in any genotype were excluded from the study (Data File S2).

**Fig. 2:**
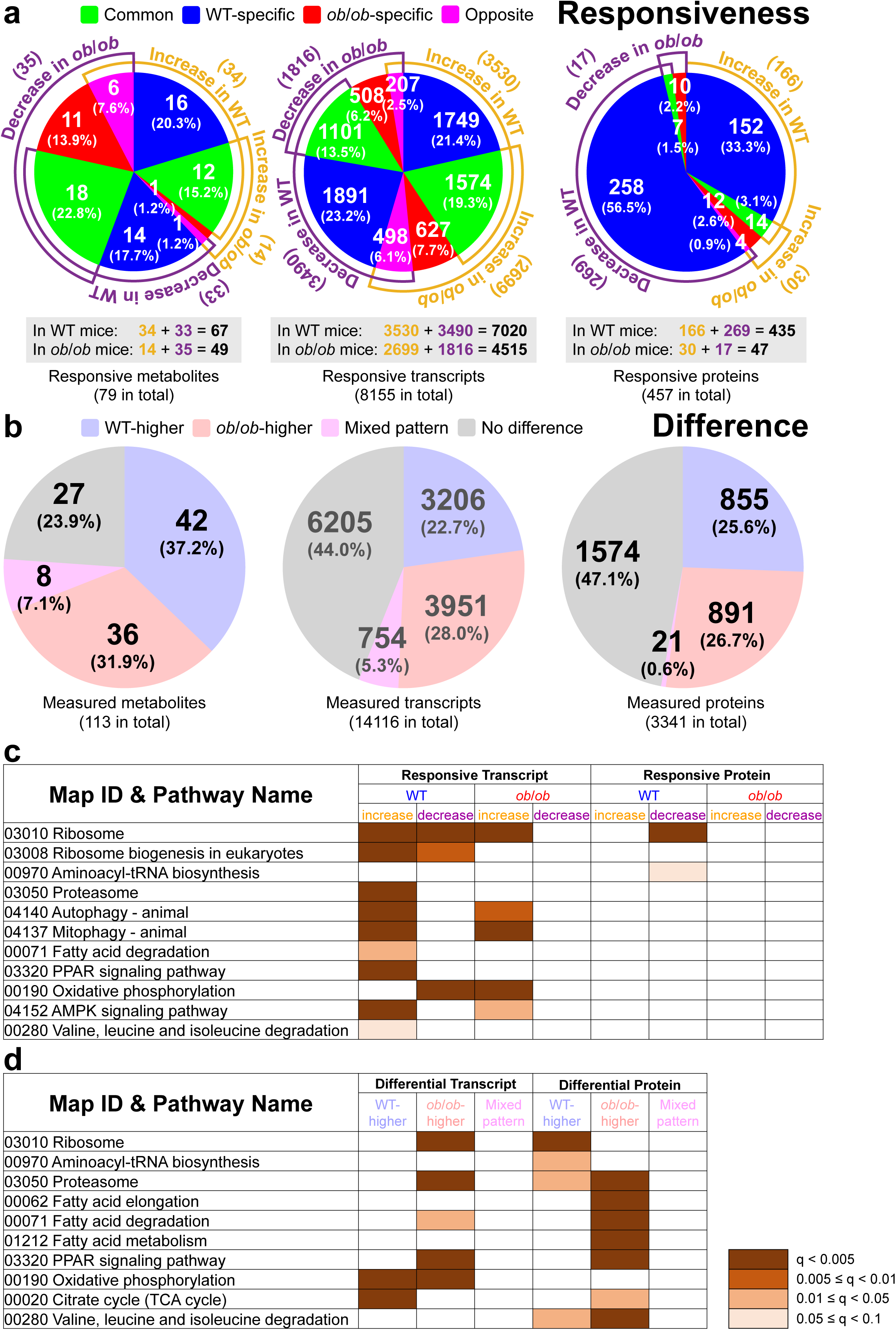
Single-omic analysis of starvation-responsive and -differential metabolites, transcripts, and proteins. (a) Pie charts showing responsiveness patterns of metabolites, transcripts, and proteins. Numbers of increased (orange) and decreased (purple) molecules are indicated in brackets around the pie charts. Below the pie charts, the total numbers of responsive molecules in WT and *ob*/*ob* mice (sum of increased and decreased molecules) are shown, with the overall number of responsive molecules (WT and *ob*/*ob* combined) provided at the bottom. Responsiveness was defined as q < 0.1, determined using one-way ANOVA with BH correction. *n* = 5 biological replicates per group. (b) Pie charts showing difference patterns of metabolites, transcripts, and proteins. Total numbers of measured molecules are indicated in brackets at the bottom. Difference was defined as q < 0.1 at any time point, determined using one-way ANOVA with BH correction. *n* = 5 biological replicates per group. (c-d) Enriched pathways in responsive (c) and differential (d) transcripts and proteins. Only selected pathways are shown. Enrichment was determined using Fisher’s exact test, with p-values adjusted for FDR using the BH method. Pathways with q < 0.1 were considered enriched.

Among starvation-responsive molecules (Fig. 2a), a greater number of responsive metabolites, transcripts, and proteins were observed in WT mice (67, 7020, and 435, respectively) compared to *ob*/*ob* mice (49, 4515, and 47, respectively). Notably, protein responsiveness was predominantly WT-specific, indicating that protein responses were largely lost in *ob*/*ob* mice.

Among starvation-differential molecules (Fig. 2b), more than half of the measured metabolites (42+36+8=86, 76.1%), transcripts (3206+3951+754=7911, 56.0%), and proteins (855+891+21=1767, 52.9%) were starvation-differential, with the similar numbers of molecules being WT-higher and *ob*/*ob*-higher.

To investigate the association of starvation-responsive -differential molecules with specific pathways, we performed KEGG^64^ (Kyoto Encyclopedia of Genes and Genomes) pathway enrichment analysis using Fisher’s exact test and applied the BH method for FDR adjustment (Fig. 2c and 2d). For metabolites, no pathways were enriched; for transcripts and proteins, enriched pathways related to protein turnover (synthesis and degradation) and lipid metabolism were selected from the full lists (Data File S3).

For responsive transcripts and proteins (Fig. 2c), the “03010 Ribosome” pathway was enriched in decreased transcripts and proteins only in WT mice, but not in *ob*/*ob* mice. The “03050 Proteasome” pathway was enriched in increased transcripts in WT mice, but not in *ob*/*ob* mice. These findings suggest that protein synthesis was inhibited at the protein level in WT mice, while protein degradation was activated at the transcript level in WT mice, but not in *ob*/*ob* mice. For differential transcripts and proteins (Fig. 2d), pathways related to fatty acid metabolism (“00062 Fatty acid elongation”, “00071 Fatty acid degradation”, “01212 Fatty acid metabolism”, and “03320 PPAR signaling pathway”) were all enriched in *ob*/*ob*-higher proteins. This result indicated elevated enzyme proteins in *ob*/*ob* mice compared to WT mice. Taken together, the above results suggest that the loss of responsiveness in protein turnover pathways and elevated enzyme proteins in lipid metabolism were part of the metabolic dysregulations in *ob*/*ob* mice compared to WT mice.

### Global trans-omics networks uncovered distinct regulatory strategies in the skeletal muscle between WT and *ob*/*ob* mice

To investigate how starvation-responsive and -differential metabolites, transcripts, and proteins regulate metabolic reactions, we constructed global trans-omics networks by integrating measured omics data with database-derived regulatory relationships (Fig. 3, see Methods). These networks consist of layers of nodes connected by edges across layers (Data File S4). The nodes represent measured molecules (metabolites, transcripts, and proteins) and reactions (metabolic and transport reactions). Nodes representing metabolic reactions are connected by molecule nodes that regulate them (fig. S2). The edges represent regulatory relationships between nodes. Edges from the “TF” layer to the “Enzyme mRNA” and “Transporter mRNA” layers represent transcriptional regulation of enzymes and transporters by transcription factors (TFs). Edges from the “Enzyme mRNA” layer to the “Enzyme Protein” layer, and from the “Transporter mRNA” layer to the “Transporter Protein” layer, represent translation processes. Edges from the “Enzyme Protein” layer to the “Metabolic Reaction” layer represent the catalytic effects of enzymes. Note that in this study, the term “enzymes” specifically refers to metabolic enzymes. Edges from the “Transporter Protein” layer to the “Transport Reaction” layer represent the facilitation of transport processes by transporters. Edges from the “Metabolite” layer to the “Metabolic Reaction” layer represent the regulation of metabolic reactions by metabolites acting as substrates/products, cofactors, allosteric activators, or allosteric inhibitors. Edges from the “Transport Reaction” layer to the “Metabolite” layer represent the impact of transport processes on metabolite levels.

**Fig. 3:**
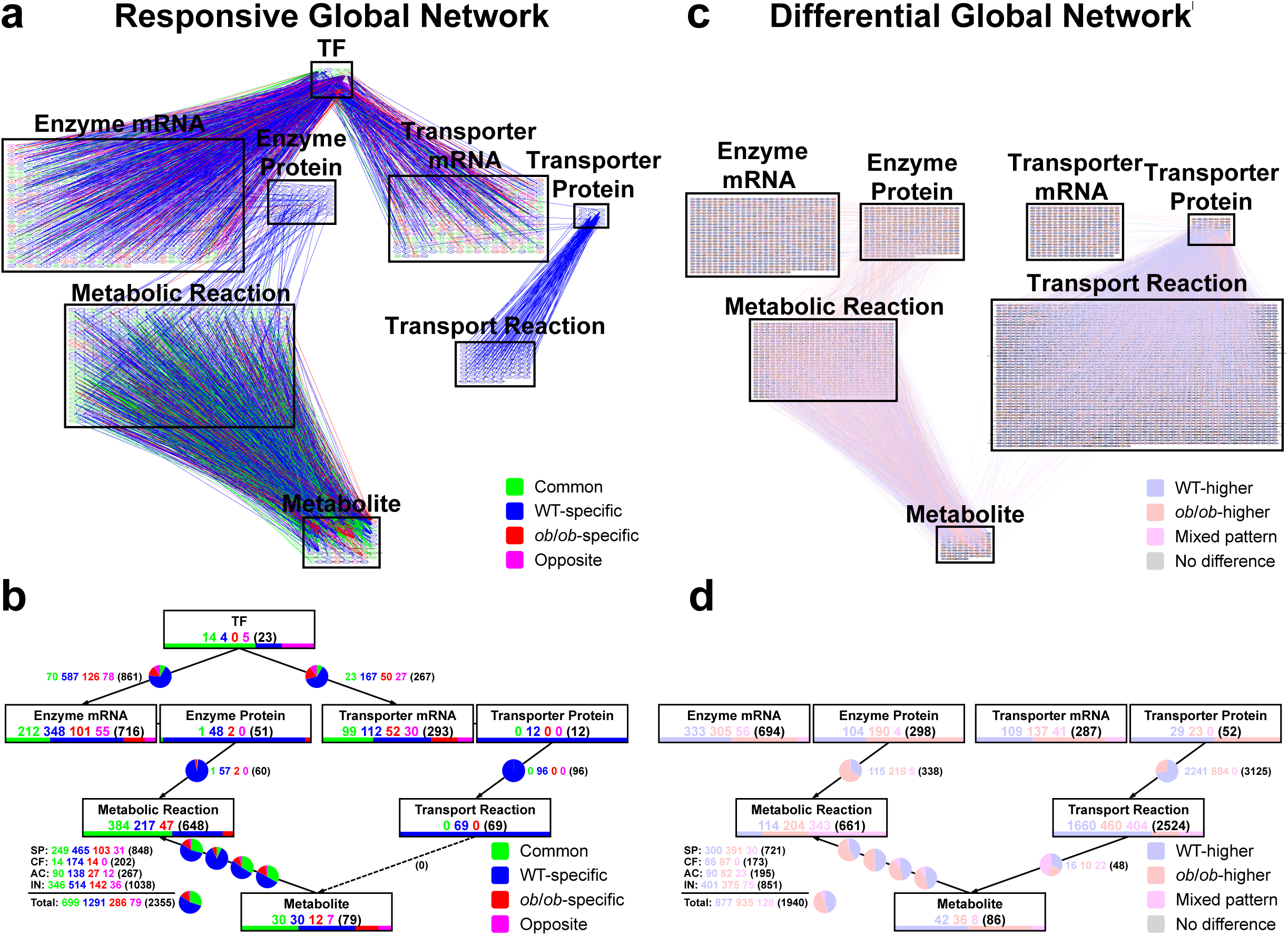
Starvation-responsive and -differential global networks. (a) Starvation-responsive global network illustrating regulatory relationships between TFs, enzymes, transporters, metabolic and transport reactions, and metabolites. (b) Calculation of responsive molecules (nodes) in each layer and responsive regulations (edges) between layers. Responsive nodes and edges were categorized as common (green), WT-specific (blue), *ob*/*ob*-specific (red), and opposite (magenta). Node numbers are shown in boxes and represented by horizontal stacked bars below each box, while edge numbers are displayed as pie charts with corresponding labels. (c) Starvation-differential global network illustrating regulatory relationships between enzymes, transporters, metabolic and transport reactions, and metabolites. (d) Calculation of differential molecules (nodes) in each layer and differential regulations (edges) between layers. Differential nodes and edges were categorized as WT-higher (light blue), *ob*/*ob*-higher (light red), and mixed pattern (light magenta). Node numbers are shown in boxes and represented by horizontal stacked bars below each box, while edge numbers are displayed as pie charts with corresponding labels. SP: Substrates/products; CF: cofactors; AC: allosteric activators; IN: allosteric inhibitors.

In the starvation-responsive global trans-omics network, only starvation-responsive molecules were included (Fig. 3a). Nodes and edges were colored according to their responsiveness, and the numbers of each category in the starvation-responsive global network were calculated (Fig. 3b). We found that molecules in the “Enzyme Protein” and “Transporter Protein” layers exhibited predominantly WT-specific responses, consistent with the responsive pattern of the entire proteome (Fig. 2a). Additionally, TF regulation of enzyme (587 out of 861, 68.2%) and transporter (167 out of 267, 62.5%) expression, as well as metabolic regulation by metabolites (1291 out of 2355, 54.8%), were also mainly WT-specific. Overall, *ob*/*ob* mice showed reduced responsiveness in enzyme and transporter proteins, as well as less regulation by responsive TFs, enzymes, transporters, and metabolites. This result indicates that WT mice use the starvation-responsive global trans-omics network to adapt to starvation, while such adaptive responses were lost in *ob*/*ob* mice.

In the starvation-differential global trans-omics network, only starvation-differential molecules were included (Fig. 3c). TFs were not included in the starvation-differential network, as TF inference is based on the responsiveness of TFs and their target genes. Nodes and edges were colored according to their difference, and the numbers of each category in the global trans-omics network in Fig. 3c were calculated (Fig. 3d). There were substantially more *ob*/*ob*-higher enzyme proteins (190) compared to WT-higher enzyme proteins (104), despite the overall proteome showing a similar number of WT-higher (855) and *ob*/*ob*-higher (891) proteins (Fig. 2b). This indicates that obesity may specifically cause elevations in enzyme protein expression, suggesting a distinct impact on metabolic regulation. This elevated enzyme protein expression suggests that *ob*/*ob* mice may attempt to adapt to starvation through chronic metabolic changes, potentially compensating for their loss of responsiveness. Taken together, WT and *ob*/*ob* mice show distinct patterns during starvation; WT mice mainly exhibit regulations by responsiveness in global network, while *ob*/*ob* mice mainly exhibit regulations by differences in global network.

### Pathway networks reveal the global loss of responsiveness and elevated enzyme proteins in the skeletal muscle of *ob*/*ob* mice

The analysis of global networks revealed that the number of regulations mediated by metabolites was substantially greater than the number of metabolites themselves. To explore the potential “key regulator metabolites” of metabolic reactions, we calculated the number of regulations mediated by each metabolite in the global network (Fig. 4a and 4b, fig. S3), and defined molecules with the top 10% highest number of regulations among all responsive/differential molecules as “key regulators”. The number of regulations counted the total number of regulations in either WT or *ob*/*ob* mice.

**Fig. 4:**
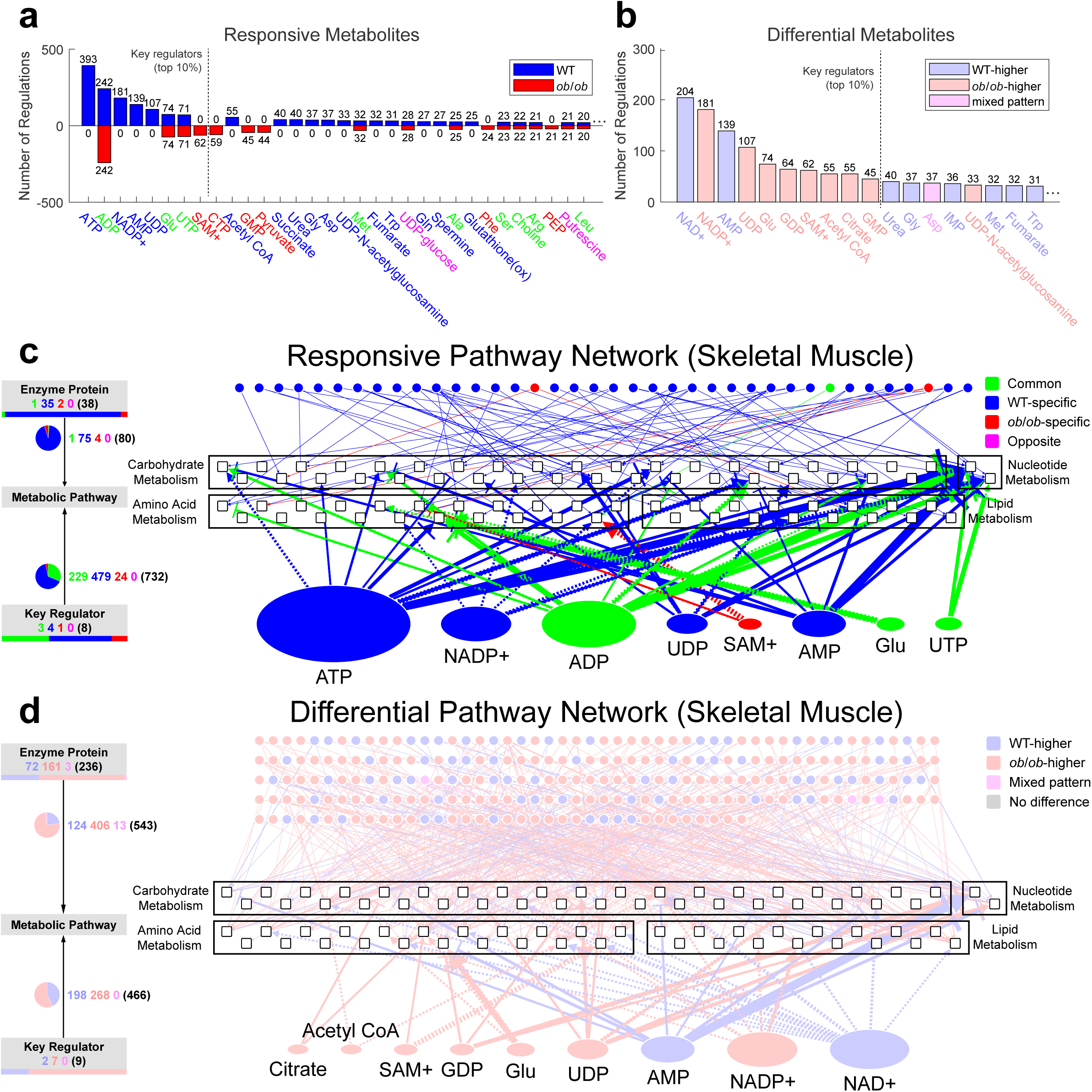
Starvation-responsive and -differential pathway networks in skeletal muscle. (a) Numbers of regulations mediated by responsive metabolites. Blue bars above zero indicate regulations in WT mice, while red bars below zero indicate regulations in *ob*/*ob* mice. Metabolite name colors denote responsiveness: common (green), WT-specific (blue), *ob*/*ob*-specific (red), and opposite (magenta). Only the top responsive metabolites are shown. (b) Numbers of regulations mediated by differential metabolites. Bar and metabolite name colors denote differences: WT-higher (light blue), *ob*/*ob*-higher (light red), and mixed pattern (light magenta). Only the top differential metabolites are shown. (c) Starvation-responsive pathway network showing the regulation of carbohydrate, amino acid, lipid, and nucleotide metabolism by responsive enzyme proteins and key regulator metabolites. The numbers of responsive enzyme proteins and key regulator metabolites are represented by the horizontal stacked bar on the left. Node sizes and edge thickness reflect the number of regulations mediated by each responsive key regulator metabolite. (d) Starvation-differential pathway network showing the regulation of carbohydrate, amino acid, lipid, and nucleotide metabolism by differential enzyme proteins and key regulator metabolites. The numbers of differential enzyme proteins and key regulator metabolites are represented by the horizontal stacked bar on the left. Node sizes and edge thickness reflect the number of regulations mediated by each differential key regulator metabolite. The names of enzyme proteins are included in Data File S9.

Among the 8 key regulators in responsive metabolites, four (ATP, NADP+, AMP, and UDP) were WT-specific, three (ADP, glutamate, and UTP) were common, and one (SAM+) was *ob*/*ob*-specific (Fig. 4a). These key regulator metabolites mediated 1207 regulations (393+242+181+139+107+74+71) in WT mice, compared to only 449 regulations (242+74+71+62) in *ob*/*ob* mice. The responsive key regulators functioning as substrates/products, cofactors, activators and inhibitors were roughly the same metabolites (fig. S3a-d). The loss of responsive key regulators in *ob*/*ob* mice caused the severe loss of regulation by metabolites observed in the starvation-responsive global network (Fig. 3b).

Among the 9 key regulators in differential metabolites, two (NAD+ and AMP) were WT-higher, seven (NADP+, UDP, glutamate, GDP, SAM+, acetyl-CoA, and citrate) were *ob*/*ob*-higher (Fig. 4b). Together, the WT-higher key regulator metabolites mediated 343 regulations (204+139), while the *ob*/*ob*-higher key metabolites mediated 598 regulations (181+107+74+64+62+55+55). The differential key regulator metabolites functioning as substrates/products, cofactors, activators and inhibitors were roughly the same metabolites (fig. S3e-h).

To further explore the regulation of metabolic reactions by responsive and differential key regulator metabolites and enzyme proteins, we condensed the global networks into starvation-responsive and -differential pathway networks (Fig. 4c and 4d), using a similar approach to our previous study^54, 55^ (see Methods). These pathway networks comprise three layers of nodes: “Enzyme Protein”, “Metabolic Pathway”, and “Key Regulator” layers. We condensed metabolic reactions into Metabolic Pathway according to our previous study.^54, 55^ Each node in the “Metabolic Pathway” represents a pathway consisting of a collection of metabolic reactions derived from the KEGG^64^ database. Nodes in the “Key Regulator” layer are the key regulator metabolites identified in Fig. 4a and 4b. Edges between layers indicate regulation of Metabolic Pathway by Enzyme Protein and Key Regulator. To provide further clarity, individual networks for carbohydrate, amino acid, lipid, and nucleotide metabolism were also plotted separately (fig. S4).

In the starvation-responsive pathway network (Fig. 4c), responsive key regulator metabolites and enzyme proteins were predominantly WT-specific, including ATP and AMP. Moreover, the regulation of metabolic pathways by key regulator metabolites and enzyme proteins were also mainly WT-specific, and was largely lost in *ob*/*ob* mice (Fig. 4c, fig. S5a and 5b). Overall, we observed the loss of responsiveness of metabolic pathway regulations by key regulator metabolites and enzyme proteins in *ob*/*ob* mice compared to WT mice in carbohydrate, amino acid, lipid, and nucleotide metabolism.

In the starvation-differential pathway network (Fig. 4d), differential key regulator metabolites and enzyme proteins were predominantly *ob*/*ob*-higher. The regulation of metabolic pathways by key regulator metabolites and enzyme proteins were also mainly *ob*/*ob*-higher, especially for enzyme proteins. Overall, we observed the metabolic pathway regulations by elevated enzyme proteins in *ob*/*ob* mice compared to WT mice in carbohydrate, amino acid, lipid, and nucleotide metabolism.

These findings indicate that the loss of responsive key regulator metabolites in *ob*/*ob* mice may underlie the observed loss of metabolite-mediated responsive regulations in *ob*/*ob* mice within the global network. Furthermore, the pathway networks elucidated the specific roles of each key regulator metabolite within the four metabolic pathways associated with energy generation.

### The global loss of responsiveness and elevated enzyme proteins were also observed in the liver of *ob*/*ob* mice

In addition to skeletal muscle, liver is another key metabolic organ. Based on our previous study of the liver of WT and *ob*/*ob* mice^29^, we re-analyzed the multi-omics data (Data File S5), identified the starvation-responsive (fig. S6a) and -differential (fig. S6b) molecules, and constructed the starvation-responsive and -differential pathway networks in the liver (Fig. 5), which displayed the similar patterns as in skeletal muscle. The key regulators (Fig. 5a and 5b) were identified by the same criteria as in skeletal muscle. Note that the identification of starvation-responsive molecules is identical to our previous study in liver^29^, but the identification of starvation-differential molecules is unique to this study. While figures similar to Fig. 5a and fig. S6a were already presented in our previous study of liver, we displayed the similar figures with minor modifications to account for differences in analytical parameters and database versions, enabling a more stringent comparison with the muscle analyses conducted in this study.

**Fig. 5:**
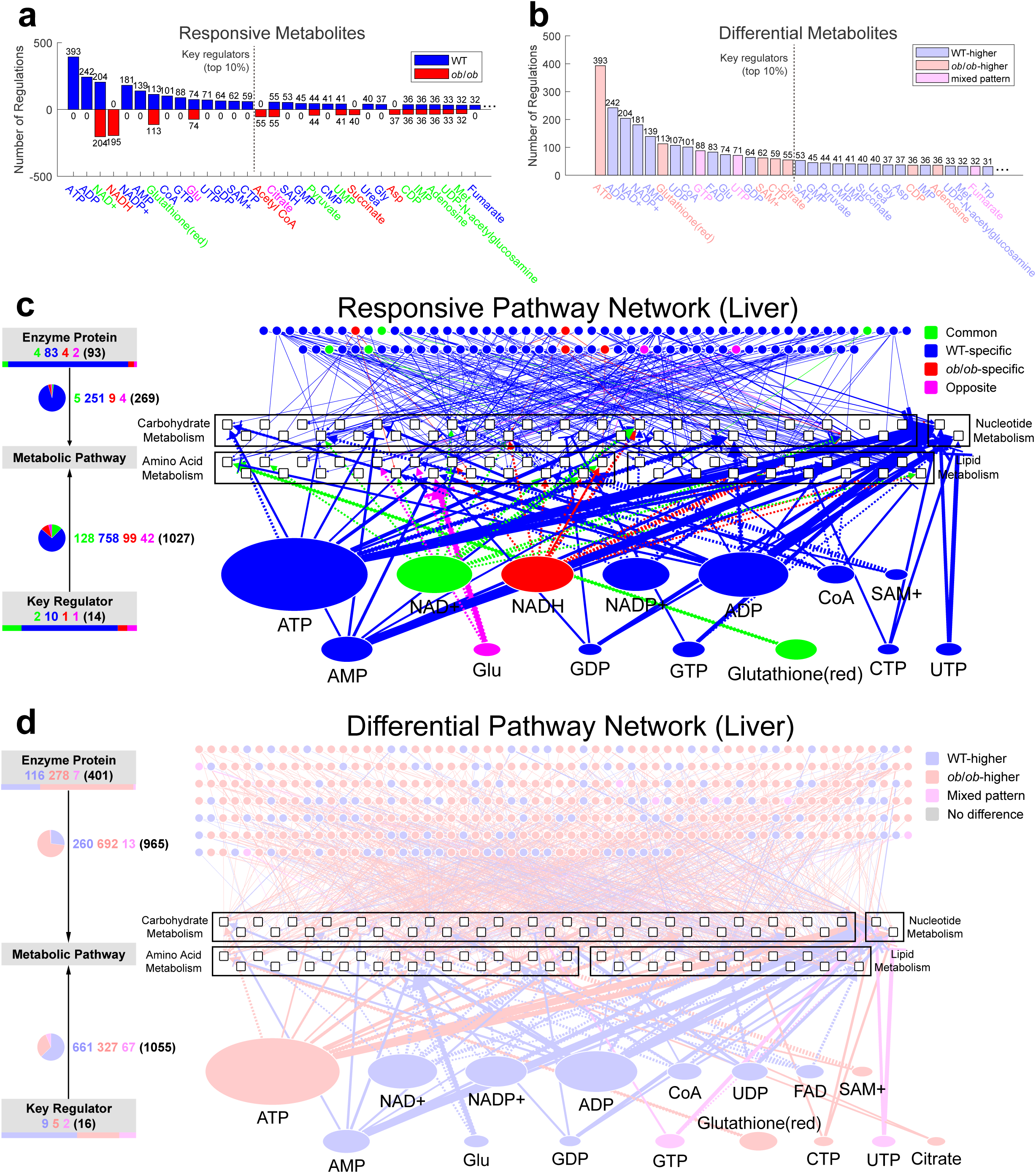
Starvation-responsive and -differential pathway networks in liver. (a) Numbers of regulations mediated by responsive metabolites. Blue bars above zero indicate regulations in WT mice, while red bars below zero indicate regulations in *ob*/*ob* mice. Metabolite name colors denote responsiveness: common (green), WT-specific (blue), *ob*/*ob*-specific (red), and opposite (magenta). Only the top responsive metabolites are shown. (b) Numbers of regulations mediated by differential metabolites. Bar and metabolite name colors denote differences: WT-higher (light blue), *ob*/*ob*-higher (light red), and mixed pattern (light magenta). Only the top differential metabolites are shown. (c) Starvation-responsive pathway network showing the regulation of carbohydrate, amino acid, lipid, and nucleotide metabolism by responsive enzyme proteins and key regulator metabolites. The numbers of responsive enzyme proteins and key regulator metabolites are represented by the horizontal stacked bar on the left. Node sizes and edge thickness reflect the number of regulations mediated by each responsive key regulator metabolite. (d) Starvation-differential pathway network showing the regulation of carbohydrate, amino acid, lipid, and nucleotide metabolism by differential enzyme proteins and key regulator metabolites. The numbers of differential enzyme proteins and key regulator metabolites are represented by the horizontal stacked bar on the left. Node sizes and edge thickness reflect the number of regulations mediated by each differential key regulator metabolite. The names of enzyme proteins are included in Data File S9.

Consistent with our previous study of liver ^29^, among the 14 key regulators in responsive metabolites (Fig. 5a), 10 (ATP, ADP, NADP+, AMP, CoA, GTP, UTP, GDP, SAM+, and CTP) were WT-specific, 2 (NAD+ and reduced glutathione) were common, one (NADH) was *ob*/*ob*-specific, and one (glutamate) was opposite. These key regulator metabolites mediated 1791 regulations (393+242+204+181+139+113+101+88+74+71+64+62+59) in WT mice, compared to only 586 regulations (204+195+113+74) in *ob*/*ob* mice.

Among the 16 key regulators in differential metabolites (Fig. 5b), 9 (ADP, NAD+, NADP+, AMP, UDP, CoA, FAD, glutamate, and GDP) were WT-higher, 5 (ATP, reduced glutathione, SAM+, CTP, and citrate) were *ob*/*ob*-higher, 2 (GTP and UTP) showed mixed pattern. Together, the WT-higher key metabolites mediated 1195 regulations (242+204+181+139+107+101+83+74+64), while the *ob*/*ob*-higher key metabolites mediated 682 regulations (393+113+62+59+55).

In the starvation-responsive pathway network (Fig. 5c), responsive key regulator metabolites and enzyme proteins were predominantly WT-specific, including ATP and AMP. Moreover, the regulation of metabolic pathways by key regulator metabolites and enzyme proteins were also mainly WT-specific, and was largely lost in *ob*/*ob* mice (fig. S7a and 7b). Overall, the loss of responsiveness of metabolic pathway regulations by key regulator metabolites and enzyme proteins was also seen in the liver of *ob*/*ob* mice compared to WT mice.

In the starvation-differential pathway network (Fig. 5d), differential key regulator metabolites were predominantly WT-higher while enzyme proteins were predominantly *ob*/*ob*-higher. The regulation of metabolic pathways by enzyme proteins were also mainly *ob*/*ob*-higher. Overall, the metabolic pathway regulations by elevated enzyme proteins in *ob*/*ob* mice compared to WT mice was also seen in the liver.

These findings indicate that, similar to skeletal muscle, the metabolic pathway regulations in liver also showed the global loss of responsiveness, accompanied by elevated enzyme proteins, in *ob*/*ob* mice during starvation.

### Loss of responsiveness in the energy-sensing pathway and disrupted downstream phosphorylation in *ob*/*ob* mice

Among the key regulator metabolites, ATP and AMP showed WT-specific responsiveness, prompting further investigation into the signaling pathways that are potentially influenced by changes in AMP and ATP levels. An energy-sensing pathway refers to a biological signaling network that monitors cellular energy status and adjusts metabolic processes to maintain energy homeostasis. A prominent example is the AMPK pathway^65, 66^, which is activated in response to a high AMP/ATP ratio, signaling low energy availability. Upon activation, AMPK promotes energy production and inhibits energy-consuming processes.^57, 58, 67, 68^ To examine the responsiveness and difference in the energy-sensing pathway in WT and *ob*/*ob* mice, we measured the phosphorylation levels of relevant proteins using western blot (Fig. 6). The phosphorylation level at each phosphosite was represented as the ratio of phosphoprotein to total protein (see Methods).

**Fig. 6:**
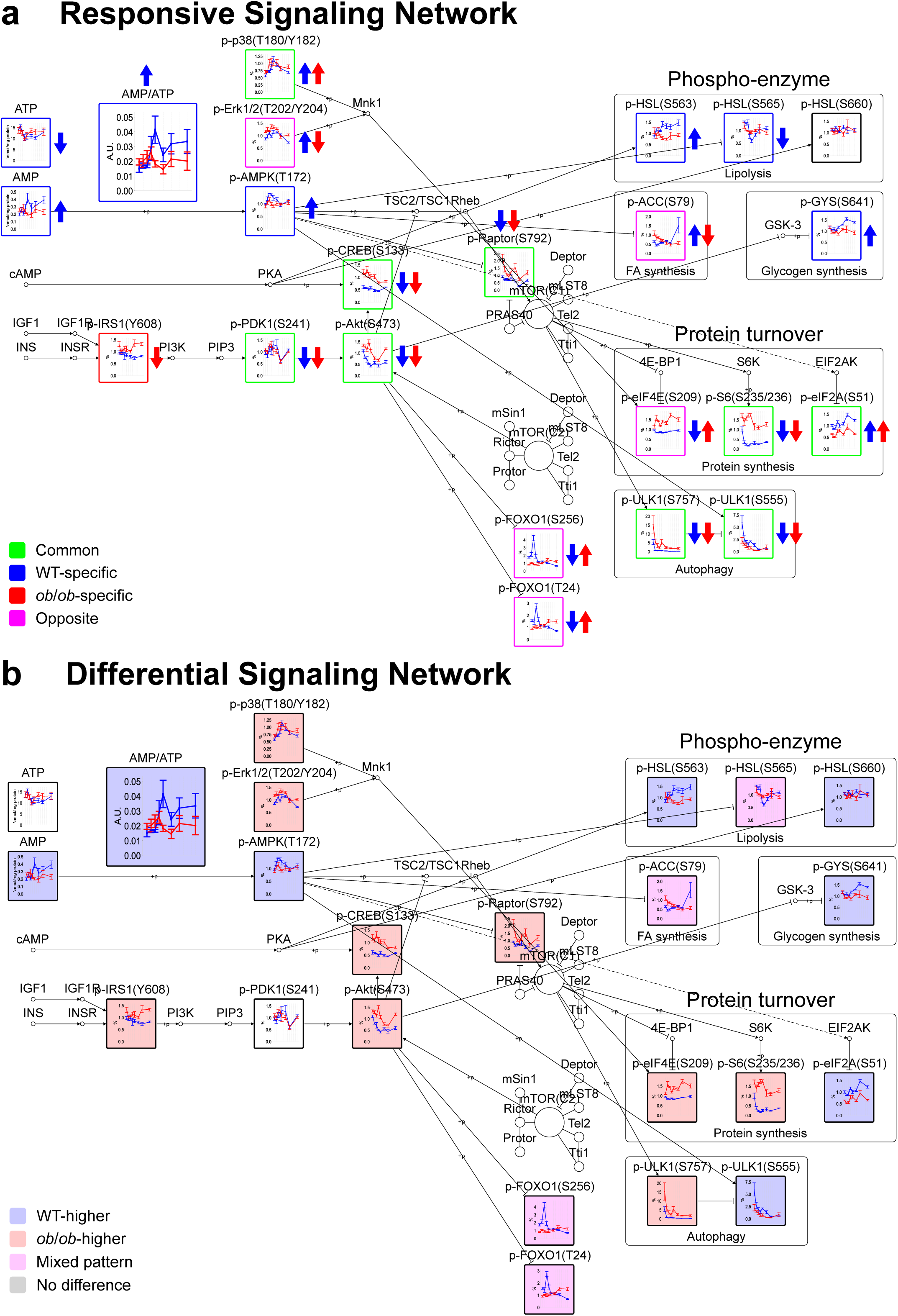
Starvation-responsive and -differential signaling networks reveal distinct regulation patterns by phosphorylation in WT and *ob*/*ob* mice (a-b) Starvation-responsive (a) and differential (b) signaling networks featuring the AMPK and PI3K/AKT/mTOR signaling pathways and their downstream proteins. Time-course data are shown as means ± SEM, with blue representing WT mice and red representing *ob*/*ob* mice. Phosphorylation events are depicted as edges connecting kinase to their target proteins, with arrowheads indicating activation and flat heads indicating inhibition. (a) Upward (increase) and downward (decrease) arrows indicate responsiveness in WT (blue) and *ob*/*ob* (red) mice, while frame colors denote the responsive patterns of phosphoproteins. (b) Fill colors indicate the differential patterns of phosphoproteins. *n* = 5 biological replicates per group.

In the starvation-responsive signaling network, AMP showed WT-specific increase while ATP showed WT-specific decrease, leading to the WT-specific increase of the AMP/ATP ratio. Accordingly, the phosphorylation of AMPK at Thr172 also increased exclusively in WT (Fig. 6a). Phosphorylated AKT, PDK1, CREB, and Raptor (the PI3K/AKT/mTOR signaling pathway) showed common decrease. In the downstream, phosphorylation of metabolic enzymes mainly showed WT-specific responsiveness. The increased phosphorylation of HSL at Ser563 and decreased inhibitory phosphorylation of HSL at Ser565 implied that lipid metabolism was possibly activated in WT mice. The opposite responses of p-ACC (Ser79) indicated that fatty acid biosynthesis may be inhibited in WT mice but activated in *ob*/*ob* mice. These results were consistent with previous findings that fatty acid utilization by skeletal muscle was impaired in obesity during postabsorptive conditions.^69-71^ In addition, glycogen synthesis was inhibited in WT mice by the WT-specific increase of p-GYS (Ser641). Previous study observed increased liver glycogen synthesis in mice fed with high-fat diet.^72^ Regulators of protein synthesis and degradation mainly showed common responses between WT and *ob*/*ob* mice. In summary, the loss of responsiveness in AMPK signaling pathway and downstream phospho-enzymes may contribute to the metabolic dysregulations of glucose and fatty acid metabolism in the skeletal muscle of *ob*/*ob* mice.

In the starvation-differential signaling network, we found that AMP level was higher in WT mice, as was the AMP/ATP ratio. As a result, p-AMPK (Thr172) also displayed higher levels in WT mice (Fig. 6b). p-IRS1, p-AKT, p-CREB, and p-Raptor (the PI3K/AKT/mTOR signaling pathway) were mainly *ob*/*ob*-higher, consistent with previous studies in obese rats^48^. In the downstream, activators (p-eIF4E^73, 74^ and p-S6^75, 76^) of protein synthesis were *ob*/*ob*-higher, while the inhibitor (p-eIF2A^77, 78^) of protein degradation was WT-higher. These results suggested impaired inhibition of protein synthesis in obese skeletal muscle during starvation. On the other hand, autophagy regulator ULK1 showed WT-higher activating phosphorylation at Ser555 and *ob*/*ob*-higher inhibitory phosphorylation at Ser757, suggesting higher activity of autophagy in WT mice^79^. Taken together, *ob*/*ob* mice showed elevated protein synthesis but reduced protein degradation in the skeletal muscle compared to WT mice. Previous studies have reported conflicting findings on protein turnover in obese subjects, including higher^80-82^, similar^83, 84^, or lower^85^ turnover compared to lean subjects. These discrepancies may be attributed to variations in experimental conditions and the animal species used. In summary, the differential phosphorylation levels of protein metabolism regulators result in distinct regulation of protein synthesis and degradation in the skeletal muscle of *ob*/*ob* mice compared to WT mice, which may contribute to the elevated enzyme proteins in the pathway networks (Fig. 4).

Similarly, in the liver of WT and *ob*/*ob* mice during starvation (fig. S8b), the AMP/ATP ratio also showed WT-specific increase, and p-AMPK (Thr172) showed a bigger increase in WT mice than in *ob*/*ob* mice, which is in principle consistent with those in skeletal muscle. Note that the responsiveness of AMP, ATP, and AMPK was already reported in our previous study of liver.^29^

These results indicate that, in the energy-sensing pathway, the responsiveness of the AMP/ATP ratio and p-AMPK was either lost or reduced in both skeletal muscle and liver of *ob*/*ob* mice during starvation.

### The global loss of responsiveness and elevated enzyme proteins represent systemic metabolic dysregulation in the skeletal muscle and liver associated with obesity

Based on the analyses of skeletal muscle during starvation, we observed a severe loss of responsiveness in metabolites and enzyme proteins in *ob*/*ob* mice (Fig. 7a and 7b). Among the key regulator metabolites, AMP and ATP were only responsive in WT mice. This led to the loss of large numbers of metabolic regulations by metabolites in *ob*/*ob* mice, as well as the failure to activate the AMPK signaling pathway and phosphorylate downstream enzyme proteins in *ob*/*ob* mice. Additionally, enzyme protein levels were consistently higher in *ob*/*ob* mice than in WT mice. In the liver, similar patterns were noted, though *ob*/*ob* mice retained a small degree of AMPK activation compared to its complete loss in skeletal muscle (Fig. 7c and 7d).

**Fig. 7:**
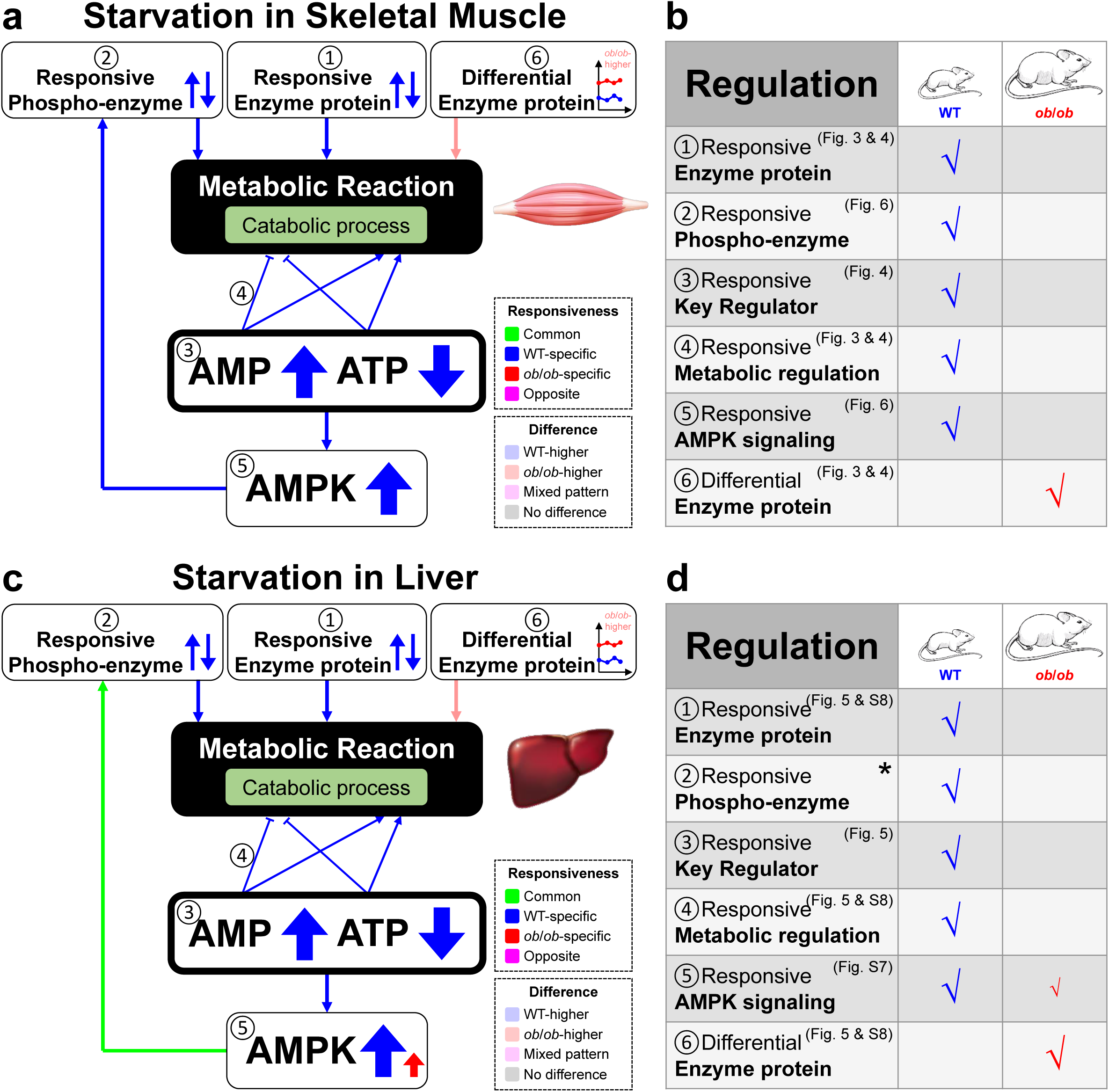
Summary of the distinct regulatory patterns in WT and *ob*/*ob* mice during starvation (a-b) Summary of key regulatory strategies in the skeletal muscle of WT and *ob*/*ob* mice during starvation. (a) Simplified network illustrating the regulation of metabolic reactions by responsive key regulator metabolites, responsive enzyme proteins, differential enzyme proteins, and responsive enzyme phosphorylation. (b) Table summarizing the primary differences between WT and *ob*/*ob* mice. (c-d) Summary of key regulatory strategies in the liver of WT and *ob*/*ob* mice during starvation. (c) Simplified network illustrating the regulation of metabolic reactions by responsive key regulator metabolites, responsive enzyme proteins, differential enzyme proteins, and responsive enzyme phosphorylation. (d) Table summarizing the primary differences between WT and *ob*/*ob* mice, with smaller check marks indicating that the regulation is present but less pronounced. Note: The asterisk (*) indicates the conclusion derived from our previous study^29^.

Taken together, the global loss of responsiveness and elevated enzyme proteins represent the systemic metabolic dysregulation associated with obesity in both skeletal muscle and liver during starvation.

## Discussion

In this study, we conducted a comprehensive trans-omics analysis to investigate the skeletal muscle adaptation to 24-hour starvation in WT and *ob*/*ob* mice. We measured the metabolome, transcriptome, proteome, and phosphoproteins (Fig. 1), and identified starvation-responsive and -differential molecules (Fig. 2). By constructing the global networks, we compared the overall regulatory patterns between WT and *ob*/*ob* mice (Fig. 3). We also identified key regulator metabolites, such as AMP and ATP, and examined their regulatory roles in the pathway networks in the skeletal muscle of WT and *ob*/*ob* mice (Fig. 4). By re-analyzing our liver data during starvation, we confirmed similar patterns in the pathway networks in the liver (Fig. 5). Furthermore, our analysis of signaling pathways revealed WT-specific activation of the AMPK signaling pathway, driven by an WT-specific increase of AMP/ATP ratio, which was absent in *ob*/*ob* mice (Fig. 6).

During adaptation to starvation, the skeletal muscle of *ob*/*ob* mice exhibited metabolic dysregulation across multiple aspects (Fig. 7). In the single-omic analysis, we observed impaired responsiveness of metabolites, transcripts, proteins in *ob*/*ob* mice, with protein responsiveness nearly entirely lost in *ob*/*ob* mice compared to WT mice. The starvation-responsive global networks and pathway networks revealed a marked reduction in the regulation of metabolic reactions by enzyme proteins and metabolites in *ob*/*ob* mice, driven by the loss of responsive molecules, particularly key regulator metabolites. Meanwhile, we observed a greater number of *ob*/*ob*-higher enzyme proteins and key regulator metabolites, suggesting distinct adaptive strategies during starvation: WT mice exhibit “responsiveness”, while *ob*/*ob* mice exhibit “difference”. In signaling networks, the activation of AMPK signaling and downstream enzyme phosphorylation were impaired in *ob*/*ob* mice, while the PI3K/AKT/mTOR signaling pathway showed abnormally higher phosphorylation levels compared to WT mice. Consequently, the phosphorylation of downstream enzymes showed inhibition of glycogen synthesis and fatty acid biosynthesis in WT mice but not in *ob*/*ob* mice, possibly indicating the failure to switch from glucose utilization to fatty acid utilization in *ob*/*ob* mice. Similar phenomenon was also noted in several previous studies. In human skeletal muscle, diminished rates of lipid oxidation during fasting conditions were observed in obesity.^46, 71, 86^ A decreased capacity to take up fatty acids has also been reported in skeletal muscle of viscerally-obese women during post-absorptive conditions.^87^ The reduced rate of lipid oxidation in obesity in previous studies suggests that the starvation adaptation by responsiveness in WT mice may be more effective than by difference in *ob*/*ob* mice.

In addition to skeletal muscle, we observed similar metabolic dysregulation during adaptation to starvation in liver. The global loss of responsiveness was observed in single-omic analysis (fig. S6), trans-omics networks (Fig. 5 and fig. S7, S9) and among molecules in energy-sensing pathway (fig. S8)).^29^ In addition, the elevated enzyme proteins were also observed in the liver of *ob*/*ob* mice (fig. S9d).

In this study, we examined the adaptation to starvation, which induces catabolic processes, driving the breakdown of stored macromolecules to generate energy and maintain metabolic homeostasis.^88-90^ In contrast, the oral glucose tolerance tests (OGTT) represents an energy influx, triggering glucose uptake and anabolic processes.^91-94^ To determine whether obesity similarly affects adaptation to energy influx, we re-analyzed our previous OGTT data measured in skeletal muscle^54^ and liver^55^ (Data File S6) and compared the findings with those observed during starvation. It should be noted that, during the re-analysis, we applied the same statistical methods and parameters as those used in this study, which differ from those in our previous studies of OGTT in the skeletal muscle and liver ^54, 55^. While the responsiveness of some molecules changed due to the different criteria, the overall patterns and conclusions remained consistent. Proteome data were not measured during OGTT in the previous studies, so we focused solely on the regulation of metabolic reactions by metabolites, excluding enzyme proteins from the analysis. In the global networks (fig. S9e and S9f), metabolic regulations by metabolites were mainly WT-specific in both skeletal muscle and liver, resembling the patterns observed during starvation. In the pathway networks (fig. S10 and S11), the loss of responsive key regulator metabolites in *ob*/*ob* mice was observed in both skeletal muscle (fig. S10) and liver (fig. S11). For energy sensors (fig. S8c and S8d), the responsiveness of AMP, ATP, and the AMP/ATP ratio was largely lost or altered in most cases in both skeletal muscle and liver. Previous studies revealed that glucose, amino acids and fatty acids that increase in the cell with the increase in food intake can inhibit AMPK.^95, 96^ Note that the pathway networks and the responsiveness of AMP and ATP were already published in our previous study in liver and skeletal muscle during OGTT.^54, 55^ However, the responsiveness presented here differs due to the use of different parameters during the re-analysis. In summary, similar metabolic dysregulation was evident during OGTT in the skeletal muscle (fig. S12a and S12b) and liver (fig. S12c and S12d) of *ob*/*ob* mice, reflecting the patterns observed during starvation. Thus, the loss of responsiveness of energy-related molecules during both OGTT and starvation represents a common feature of the anabolic and catabolic metabolic dysregulations associated with obesity.

In addition to skeletal muscle and liver, adipose tissue is another key metabolic organ. In the previous study of rats, the AMP/ATP ratio and AMPK phosphorylation levels increased in adipose tissue after 15 hours of fasting^97^, indicating that the changes in AMP, ATP and AMPK were the common adaptive responses across multiple energy-regulating tissues including skeletal muscle, liver and adipose tissue during starvation. These changes were lost in the skeletal muscle and liver of *ob*/*ob* mice; however, response of AMP, ATP and AMPK in adipose tissue of *ob*/*ob* mice has thus far not been investigated yet, which will be examined in a separate study.

One of the limitations of this study is that our TF inference did not include an important TF, FOXO1, because ChIP-Atlas^98^ lacks experimental data for FOXO1 in the skeletal muscle of mice. Although we considered using alternative sources to derive the target genes of FOXO1, doing so would compromise the comparability across TFs, potentially biasing the analysis. With the future expansion of ChIP-Atlas^98^ data to include FOXO1 in skeletal muscle, future studies will be able to integrate this crucial TF into trans-omics analyses more comprehensively. This expansion would enable a more detailed understanding of FOXO1’s regulatory role in metabolic adaptations to starvation. Meanwhile, the included TFs may include circadian TFs, such as *Nr1d1*^99^, which exhibit increase or decrease over the 24-hour period and could be misinterpreted as starvation-responsive.

Another limitation is that we only used *ob*/*ob* mice to model obesity, which cannot exclude the effect of leptin deficiency specific to this genetic model. Previous studies have found that leptin can selectively stimulates phosphorylation and activation of AMPK in skeletal muscle^100^, which occurrs independently of energy-related metabolites^101^. These findings suggest that while *ob*/*ob* mice can provide insights, they may not fully represent the broader spectrum of obesity. Future studies may consider to include other obesity models, such as diet-induced obesity (DIO) mice. Additional models will help to differentiate between the effects of leptin deficiency and other mechanisms driving obesity. Comparative studies using these models can provide a more comprehensive understanding of the metabolic adaptations to starvation across different types of obesity.

## Materials and Methods

### Mouse experiments

All the animal experiments were conducted after approval by the animal ethics committee of the University of Tokyo. Animal experiments were conducted as previously described^29^ and muscle was collected from the identical mice of the study. Precisely, ten-week-old C57BL/6 WT and *ob*/*ob* male mice were obtained from Japan SLC Inc. Starvation started at approximately 18:00, and mice were subjected to starvation for 0, 2, 4, 6, 8, 12, 16, or 24 hours. At the corresponding time points, the mice were euthanized by cervical dislocation, and gastrocnemius and soleus muscles were dissected and immediately frozen in liquid nitrogen. The frozen muscle was pulverized with ShakeMaster NEO (BMS) followed by multi-omics measurements.

For the measurement of blood insulin and blood glucose, a different set of WT and *ob*/*ob* mice were starved for 0, 2, 4, 6, 8, 12, 16, 24 hours. At each time point, blood glucose was measured using Lab Gluco (RIJ, 4239R1006) and blood was collected in the presence of aprotinin (A1153-5MG, SIGMA). Plasma was collected and the insulin concentration was determined using Insulin ELISA Kit (Wako). U-type (633-03411) was used for WT mice and T-type (634-01481) was used for *ob*/*ob* mice.

### Metabolomic analysis

Metabolomic analysis was conducted as previously described^55^. For muscle, total metabolites and total proteins were extracted with methanol:chroloform:water (2.5:2.5:1). Approximately 40 mg of muscle was homogenized in 500 µl of ice-cold methanol containing internal standards [20 μM l-methionine sulfone (Wako), 2-morpholinoethanesulfonic acid, monohydrate (Dojindo), and d-camphor-10-sulfonic acid (Wako)] to normalize peak intensities of mass spectrometry (MS) among runs. Then, 500 μL of chloroform and 200 μL of water were applied and mixed. The mixture was centrifuged at 4,600 × *g* for 15 min at 4°C, the aqueous layer was then filtered through a 5 kDa cutoff filter (UFC3LCCNB-HMT, Human Metabolome Technologies) to exclude protein contamination.

Proteins were extracted from the interphase and organic layers by adding 800 μL of ice-cold methanol followed by centrifugation at 12,000 × *g* for 15 min at 4°C. The pellet was then washed using 1 mL of ice-cold 80% (v/v) methanol, resuspended in 500 μL water, and sonicated with Bioruptor (UCW-310, COSMO BIO). The pellet was lysed by adding 500 μL of 2% SDS and 100 mM Tris-HCl pH8.8 for 1 hour at 4LJ with gentle mixing. The samples were sonicated using Handy Sonic (UR-20P, TOMY). Total protein concentration was determined using the bicinchoninic acid (BCA) assay (23227, Thermo Fisher Scientific) and utilized for normalizing concentration among samples.

The filtrate was lyophilized and re-suspended in 50 μL of water containing the reference compounds [200 μM each of trimesate (206-03641, Wako) and 3-aminopyrrolidine (404624, Sigma-Aldrich)]. Quantification of metabolites was conducted using an Agilent 1600 Capillary Electrophoresis system (Agilent technologies), a G1603A Agilent CE-MS adapter kit, and a G1607A Agilent CE electrospray ionization (ESI)–MS sprayer kit. A fused silica capillary [50 μm internal diameter (i.d.) × 100 cm] was used with 1 M formic acid as the electrolyte^102^ for cationic compounds. Sheath liquid (methanol/water (50%, v/v) containing 0.01 μM hexakis (2,2-difluoroethoxy) phosphazene) was delivered at 10 μl/min. ESI–time-of-flight (TOF) MS was performed in positive ion mode, with the capillary voltage set to 4 kV. The acquired spectrum was each recalibrated automatically based on the masses of the reference standards [^13^C isotopic ion of a protonated methanol dimer (2 MeOH + H)]^+^, mass/charge ratio (*m*/*z*) 66.0631 and [hexakis(2,2-difluoroethoxy)phosphazene + H]^+^, *m*/*z* 622.0290. To identify metabolites, relative times of migration of all peaks to the reference compound 3-aminopyrrolidine were calculated, and their *m*/*z* values and relative migration times to the metabolite standards were compared. To quantify the metabolites, peak areas of each metabolite was compared to calibration curves generated by methionine sulfone as an internal standard. The other parameters and conditions were the same as previously described.^103^ COSMO (+) (chemically coated with cationic polymer) capillary (50 μm i.d. by 105 cm) (Nacalai Tesque, Kyoto, Japan) was used with a 50 mM ammonium acetate solution (pH 8.5) as the electrolyte for anionic metabolites. Sheath liquid (methanol/5 mM ammonium acetate (50%, v/v) containing 0.01 μM hexakis (2,2-difluoroethoxy) phosphazene) was delivered at 10 μl/min. ESI-TOFMS was performed in negative ion mode, and the capillary voltage was set to 3.5 kV. For the reference and the internal standards, trimesate and d-camphor-10-sulfonic acid were used, respectively. The other parameters and conditions were same as described previously.^104^ The Agilent MassHunter software (Agilent technologies) was used for data analysis.^102-104^

F1,6P and F2,6P were measured separately using IC-QEMS^105^. To separate metabolite, a Dionex IonPac AS11-HC-4 µm column (250 × 0.4 mm, 4 µm; Thermo Fisher Scientific) was used at 35°C. KOH was used as an eluent at the speed of 0.02 mL/min. The gradient was: 1 mM from 0 to 2 min, 20 mM at 16 min, 100 mM at 35 min. Isopropanol containing 0.1% acetic acid was used as a sheath solution at the speed of 5 µl/min. MS was conducted in the ESI negative-ion mode. The parameters were as follows: sheath gas, 20 (arbitrary units); auxiliary gas, 10 (arbitrary units); spray voltage, 4.0 kV; S-lens, 50 (arbitrary units); capillary temperature, 300°C. Data acquisition was performed in full MS scan mode. Parameters of the scanning were: resolution, 70,000; auto-gain control target, 3×10^6^; maximum ion injection time, 100 ms; scan range, 70–1,000 *m*/*z*.

### Transcriptomic analysis

Total RNAs were extracted from muscle using Maxwell ® RSC Tissue simplyRNA Purification Kit (AS1340, Promega). The tissue was homogenized in 200 μl of homogenization buffer containing 1-thioglycerol, followed by lysis by addition of 200 μl of lysis buffer and 30 μl of Proteinase K solution. After 10 minutes incubation, all the lysate was applied to the cartridge of Maxwell and total RNA was extracted. The amount and quality of RNAs were evaluated by Nanodrop (Thermo Fisher Scientific) and the 2100 Bioanalyzer (Agilent Technologies). The cDNA libraries were obtained using the TruSeq Stranded mRNA Kit (Illumina, San Diego, CA, USA). The acquired cDNAs were then sujected to 150 base pair (bp) paired-end sequencing on an Illumina NovaSeq6000 Platform (Illumina). The sequences were then aligned to the reference genome of mice obtained from the Ensembl database^106, 107^ (GRCm38/mm10, Ensembl release 97) using the STAR software package (v.2.5.3a)^108^. The RSEM tool^109^ (v.1.3.0) was utilized to estimate gene expression levels from the aligned sequences. The obtained read counts were used as inputs for edgeR software^61^. For other analyses, transcripts per kilobase million (TPM) were used.

### Proteome analysis

The muscle samples were added 0.1% trifluoroacetic acid (TFA) in acetonitrile (ACN) and a 5 mm zirconia bead (TOMY SEIKO, Tokyo Japan), and crushed using a Tissue Lyser (Qiagen, Hilden, Germany) to removal of the bead. After removal of the supernatant by centrifugation at 15,000 × g for 15 min at 4°C, the precipitates were dissolved in 0.5% sodium deoxycholate in 100 mM Tris-HCl pH 8.5 by using a water bath-type sonicator (Bioruptor II, CosmoBio, Tokyo Japan). Protein concentration in the protein extracts was determined using a BCA protein assay kit (Thermo Fisher Scientific, Waltham, MA, USA) and adjusted to 1 µg/μL with 0.5% sodium deoxycholate in 100 mM Tris-HCl pH 8.5. 20 µL of 1 µg /μL protein extracts were incubated at 50°C for 30 minutes in the presence of 10 mM DTT, followed by alkylation with 30 mM iodoacetamide for 30 min at room temperature in the dark. To stop the iodoacetamide reaction, the alkylated sample was incubated at room temperature for 10 minutes with 60 mM cysteine. After adding 150 μL of 50 mM ammonium hydrogen carbonate, the protein suspension was then subjected to digestion with 400 ng Lys-C and 400 ng trypsin for overnight at 37°C. The digested sample was acidified with 30 μL of 5% TFA and then sonicated using Bioruptor II. The resultant sample was desalted using an SDB-STAGE tip (GL Sciences, Tokyo, Japan) according to the manufacturer’s protocol, and was dried in a centrifugal evaporator (miVac Duo concentrator, Genevac, Ipswich, UK) and redissolving in 3% ACN in 0.1% formic acid. The peptide concentration was determined with the BCA protein assay kit (Thermo Fisher Scientific) and adjusted to 200 ng/μL with 3% ACN in 0.1% formic acid, prior to MS analysis.

600 ng of digested peptides was directly injected onto an Aurora column (C18, 75 μm ID, 25 cm length, 1.6 μm beads, IonOpticks, Victoria, Australia) at 60°C and then separated with a 80-min gradient (A = 0.1% formic acid in water, B = 0.1% formic acid in 80% ACN) consisting of 0 min 5% B, 70 min 35 %, 80 min 60% B at a flow rate of 200 nL/min using an UltiMate 3000 RSLCnano LC system (Thermo Fisher Scientific). The peptides eluted from the column were analyzed by overlapping window DIA-MS (PMID: 30671891, 35522919) using an Orbitrap Exploris 480 (Thermo Fisher Scientific) with an InSpIon system (AMR, Tokyo, Japan) (PMID: 37036810). MS1 spectra were collected in the range of *m/z* 495–745 at a 15,000 resolution to set an AGC targets of 3 × 10^6^ and a maximum injection time of “Auto”. MS2 spectra were collected at *m/z* 200-1,800 at a 30,000 resolution to set an AGC targets of 3 × 10^6^, a maximum injection time of “Auto”, and stepped normalized collision energies of 22, 26 and 30%. The isolation width for MS2 was set to 4 Th, overlapping window patterns at *m/z* 500–740 were used for window placements optimized via Skyline 4.1 (PMID: 20147306). The compensation voltage value and outer/inner temperatures on FAIMS were −45 V and 90 °C/100 °C, respectively.

MS files were searched against a mouse spectral library using Scaffold DIA v3.0.1 (Proteome Software, Inc., Portland, OR). The mouse spectral library was generated from Ensembl ver. 97 database by Prosit (PMID: 32214105, PMID: 31133760). The Scaffold DIA search parameters were set as: experimental data search enzyme, trypsin; maximum missed cleavage sites, 1; precursor mass tolerance, 10 ppm; fragment mass tolerance, 10 ppm; static modification, cysteine carbamidomethylation. The protein identification threshold was set both peptide and protein false discovery rates of less than 1%. Peptide quantification was computed by EncyclopeDIA algorithm v1.2.2 (PMID: 30510204) in Scaffold DIA. For each peptide, the four highest quality fragment ions were used for quantitation. The protein quantification was estimated from the summed unique peptide quantification.

### Western blot

The expression and phosphorylation of signaling molecules and their downstream target proteins were detected by western blotting on the muscle lysates (fig. S13). α-Tubulin was also measured to represent total protein content in each sample. Total proteins were extracted during sample preparation of metabolomic analysis with methanol:chloroform:water (2.5:2.5:1) extraction. The lysate was boiled in sample buffer (91 mM Tris-HCl pH6.8, 3.48% SDS, 4.7% glycerol, 2% 2-mercaptoethanol, 0.00094% BPB), separated on SDS–polyacrylamide gel electrophoresis (PAGE) and blotted with the antibodies (Data File S7). Internal control (IC) was made by mixing equal amounts of all the 80 samples (8 time points × 2 genotypes × 5 replicates = 80 samples). Signals were detected using the Immobilon Western Chemiluminescent HRP Substrate (Millipore), and measured using a luminoimage analyzer (Fusion System Solo 7S; M&S Instruments Inc., Osaka, Japan) and quantified using Fiji software (ImageJ; National Institutes of Health, Bethesda, MD, USA)^110^. Quantification of protein bands from western blot membranes was performed using Fiji software. For each membrane, the raw data of 40 samples was divided by the arithmetic mean of the two ICs, and then normalized by the α-Tubulin level in each sample. The normalized data of phosphoproteins was divided by the normalized data of total proteins to represent the phosphorylation level of that protein (at that phosphosite). In the subsequent sections of the article, phosphorylation level refers to this phosphoprotein/total protein ratio.

### Identification of starvation-responsive and -differential molecules

To identify starvation-responsive molecules in the metabolome, transcriptome, proteome, and phosphoprotein data, one-way ANOVA (ANOVA-like testing of edgeR for the transcriptome) was performed to compare different time points in each genotype. Benjamini-Hochberg (BH) method was applied for false discovery rate (FDR) adjustment. Molecules with q values less than 0.1 were defined as starvation-responsive. Based on the area under the curve (AUC) of the log2-normalized time courses, molecules with positive AUCs were classified as increased, while those with negative AUCs were classified as decreased. The between-group degrees of freedom are 7; while the within-group degrees of freedom are 32.

To identify starvation-differential molecules in the metabolome, transcriptome, proteome, and phosphoprotein data, one-way ANOVA (ANOVA-like testing of edgeR for the transcriptome) was performed to compare WT and *ob*/*ob* mice at each time point. Benjamini-Hochberg (BH) method was applied for false discovery rate (FDR) adjustment. Molecules with q values less than 0.1 at any time point were defined as starvation-differential. WT-higher molecules were those that were higher in WT mice or similar between the two genotypes across all time points. Similarly, *ob*/*ob*-higher molecules were either higher in *ob*/*ob* mice or comparable between genotypes at all time points. Molecules showing a mixed pattern were defined as being higher in WT mice at certain time points and higher in *ob*/*ob* mice at others. The between-group degrees of freedom are 1; while the within-group degrees of freedom are 9.

### TF Inference

TF inference was based on ChIP-Atlas^98^ data derived from mouse muscle tissue (mm10, Muscle) to determine target gene information. TF binding to target genes was affirmed if the binding scores exceeded 50 in half or more than half of the experiments. GO annotations^111, 112^ from UniProt^113^ were used to ascertain TFs with DNA-binding transcription factor activity (GO:0003700) or RNA polymerase II-specific DNA-binding transcription factor activity (GO:0000981). General TFs and coTFs (coactivators and corepressors) lacking strong sequence-specificity were excluded. Henceforth, “TFs” refers exclusively to DNA-binding transcription factors. TFs with “DNA-binding transcription activator activity” (GO:0001216) and “DNA-binding transcription activator activity, RNA polymerase II-specific” (GO:0001228) were deemed as activators, while those with “DNA-binding transcription repressor activity” (GO:0001217) and “DNA-binding transcription repressor activity, RNA polymerase II-specific” (GO:0001227) were deemed as repressors; some exhibited both functions. TF regulation of target genes during starvation was inferred using ChIP-Atlas^98^ binding data, GO annotations^111, 112^, and responsiveness patterns of TFs and target genes. The TF regulation was confirmed by transcriptome data of TF and target genes, and whether the TF is an activator or a repressor. For each TF-target gene binding pair, if the TF and target gene showed concordant responsiveness (both increase or both decrease) and the TF possessed activator activity, the regulation was confirmed. Conversely, if the responsiveness was discordant and the TF possessed repressor activity, the regulation was also confirmed (Data File S8). In this step, the transcriptome data for TFs were used.

### Pathway enrichment analysis

Pathway enrichment analysis was conducted separately for increased and decreased transcripts and proteins, as well as for WT-higher, *ob*/*ob*-higher, and mixed pattern transcripts and proteins. Gene lists associated with each pathway were obtained from the KEGG^64^ database (accessed on 2023/06/13). Fisher’s exact test was applied to calculate p-values, which were further adjusted for FDR using the BH method to obtain q-values. Pathways with q-values below 0.1 were identified as enriched.

### Construction of starvation-responsive and -differential global networks

The information of substrates, products, and enzymes participating in each metabolic reaction was derived from the KEGG^64^ database. The pathways maps were also obtained from the KEGG^64^ database. Allosteric regulators (activators and inhibitors) and cofactors for each enzyme of the metabolic reactions were obtained from the BRENDA^114^ database. Transcription factors for each gene were inferred using the ChIP-Atlas^98^ database and GO annotations^111, 112^ without specifying whether they are activators or repressors. Transporter information was based on the Transporter Classification Database (TCDB)^115^. Given that TCDB includes transporters from various species, including non-mouse orthologs, bioDBnet^116^ was employed to convert transporter orthologues to their respective mouse versions, where such mappings were available. The multi-layered trans-omics network was constructed using VANTED software (V2.6.5)^117^.

The network consisted of layers of molecules (nodes) with regulatory connections (edges) between them. TFs were considered to regulate the expression of enzymes and transporters. Enzyme proteins regulated metabolic reactions, while transporter proteins regulated transport reactions. Metabolites regulated metabolic reactions by functioning as substrates/products, cofactors, and allosteric regulators (activators and inhibitors), and were, in turn, regulated by transport reactions. Transport reactions facilitate the exchange of metabolites between skeletal muscle and blood.

Nodes represent measured molecules (including metabolites, transcripts and proteins) and reactions (including metabolic reactions and transport reactions). In starvation-responsive networks, the frame colors of nodes indicate their responsiveness in WT and *ob*/*ob* mice. In networks of both genotypes, blue indicates WT-specific response; red indicates *ob*/*ob*-specific response; green indicates common response (increase or decrease in both genotypes); magenta indicates opposite response (increase in one genotype and decrease in the other). The responsiveness of reactions was determined based on whether they were regulated only in WT mice (blue), only in *ob*/*ob* mice (red), or both (green). In starvation-differential networks, the fill colors of nodes indicate their difference between WT and *ob*/*ob* mice: light blue indicates WT-higher, light red indicates *ob*/*ob*-higher, light magenta indicates mixed pattern. The difference of reactions was determined based on their regulators.

Edges represented regulatory connections between nodes, categorized into two types: activating/inhibitory regulations (solid lines) and biochemical components (dotted lines). Activating regulations were indicated by pointed arrowheads, while inhibitory regulations were indicated by flathead arrows. Biochemical components included substrates or products of metabolic reactions, as well as cofactors necessary for enzyme-catalyzed metabolic reactions. The responsiveness and difference of a regulation was determined by the regulator, with the same color definition as nodes. Note that for TF regulations, the responsiveness of regulation is not necessarily the same as the TF, because TF regulations were inferred based on the responsiveness of both the TF and the target gene, not by the TF alone.

### Construction of starvation-responsive and -differential pathway networks

The starvation-responsive and -differential pathway networks were derived from the starvation-responsive and -differential global networks using the same information. The metabolic reactions belonging to pathways associated with carbohydrate (“1.1 Carbohydrate metabolism” and “1.7 Glycan biosynthesis and metabolism”), amino acid (“1.5 Amino acid metabolism” and “1.6 Metabolism of other amino acids”), lipid (“1.3 Lipid metabolism”), and nucleotide metabolism (“1.4 Nucleotide metabolism”) were included in the pathway networks, with the information of pathways from KEGG^64^ database. In the pathway networks, the regulations of all metabolic reactions in each pathway were added together. Metabolites that ranked within the top 10% for the highest number of regulations in responsive or differential metabolites were considered key regulators and included in the networks, with node size and edge thickness reflecting the number of regulations. Edges representing less than 5 regulations were omitted from the networks. Substrate/product, cofactor, activator, and inhibitor regulations were calculated independently. All regulations by enzyme proteins were included.

In the separated pathway networks for carbohydrate, amino acid, lipid, and nucleotide metabolism, the time courses of each measured molecule were plotted inside the nodes. The node sizes of key regulators were set to the same for easier visualization.

### Construction of starvation-responsive and -differential signaling network

The information on signaling network was obtained from KEGG^64^ database and publications. For phosphoproteins, the ratio of phosphorylated proteins to total proteins was used to determine the phosphorylation level. Phosphorylation at different phosphosites on the same protein was considered as different molecules, and was plotted separately.

## Supporting information

Legends for Supplementary Figures

## Data Availability

The RNA-seq data generated during the current study are available in the DDBJ^118^BioProject repository with links to BioProject accession number PRJDB19859. The mass spectrometry proteomics data generated during the current study are available in the jPOST repository^119^ with links to accession number JPST003499.

## Code Availability

The code used for the analysis in this paper is available from corresponding author upon request.

## Acknowledgments

We thank Maki Ohishi and Ayano Ueno (Keio University) for their expertise and assistance with metabolome analysis using CE-MS. We also thank Kazusa DNA Research Institute for conducting the proteomic measurements. We also thank our laboratory members for critically reading this manuscript and technical assistance with the experiments. The computational analysis of this work was performed in part with support of the supercomputer system of the National Institute of Genetics (NIG), Research Organization of Information and Systems (ROIS).

This study was supported by the Japan Society for the Promotion of Science (JSPS) KAKENHI grant numbers JP17H06299, JP17H06300, JP18H03979, JP21H04759, JP23H04939, and JP23H04946 to S.K., and by Japan Science and Technology Agency (JST) as part of CREST (JPMJCR2123 to S.K., Y. Inaba, and T.S.) and of Adopting Sustainable Partnerships for Innovative Research Ecosystem (ASPIRE), Grant Number JPMJAP24B1; The Uehara Memorial Foundation (to S.K.); K.M. receives funding from a Grant-in-Aid for Early-Career Scientists (JP21K15342). T.K. receives funding from a Grant-in-Aid for Early-Career Scientists (JP21K16349). A.Hatano receives funding from a Grant-in-Aid for Early-Career Scientists (JP22K15034). This work was also supported by the Japan Society for the Promotion of Science (JSPS) KAKENHI grant number JP22H04925 (PAGS); in part by the MEXT Cooperative Research Project Program, Medical Research Center Initiative for High Depth Omics, and CURE:JPMXP1323015486 for MIB, Kyushu University; and AMED Grant Number JP21zf0127001 (T.S.), JST, MEXT KAKENHI Grant Number JP23H04946 (T.S.) and World Premier International Research Center Initiative (WPI), Human Biology-Microbiome-Quantum Research Center (Bio2Q) (T.S.), MEXT, Japan.; and JST FOREST Program (Grant Number JPMJFR2052, to A. Hirayama). T.T. receives funding from a Grant-in-Aid for Early-Career Scientists (JP20K19915). H.O. receives funding from a Grant-in-Aid for Early-Career Scientists (JP22K17992). S.O. receives funding from a Grant-in-Aid for Early-Career Scientists (JP21K14467). Y. Inaba also receives AMED-PRIME (JP23gm6910002).

## Author contributions

D.L., A. Hatano, S.O., K.M., and S.K. designed the project. A. Hatano performed animal experiments. A. Hatano and K.M. prepared samples for omic measurement. T.S. and A. Hirayama performed metabolomic analysis using CE-MS and IC-QEMS. Y.S. performed transcriptomic analysis using RNA-seq. D.L., T.T., and H.O. analyzed the RNA-seq data. M.M and A. Hatano performed proteomic analysis using MS. D.L. performed the western blot analysis. D.L. analyzed the omics data. D.L. performed the visualization of the results. H.I. and Y.I. helped with the biological interpretation. D.L. and S.K. wrote the manuscript. All authors read and approved the final manuscript.

---

**fig. S1.**
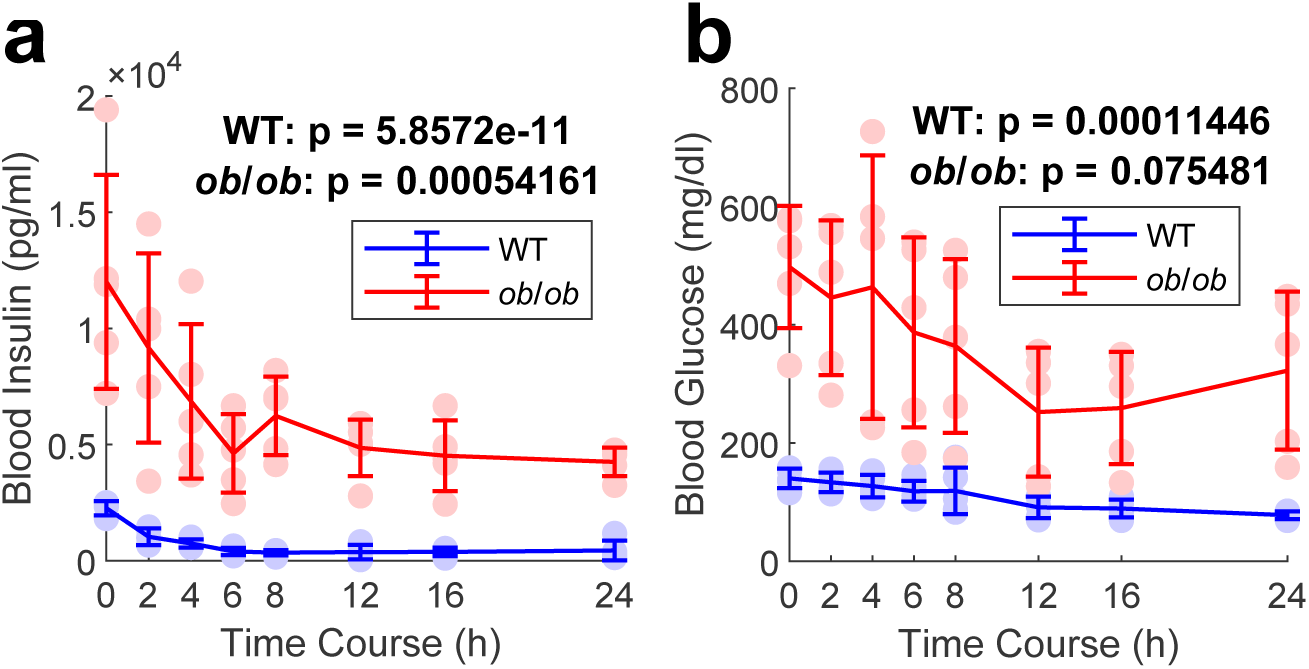

**fig. S2.**
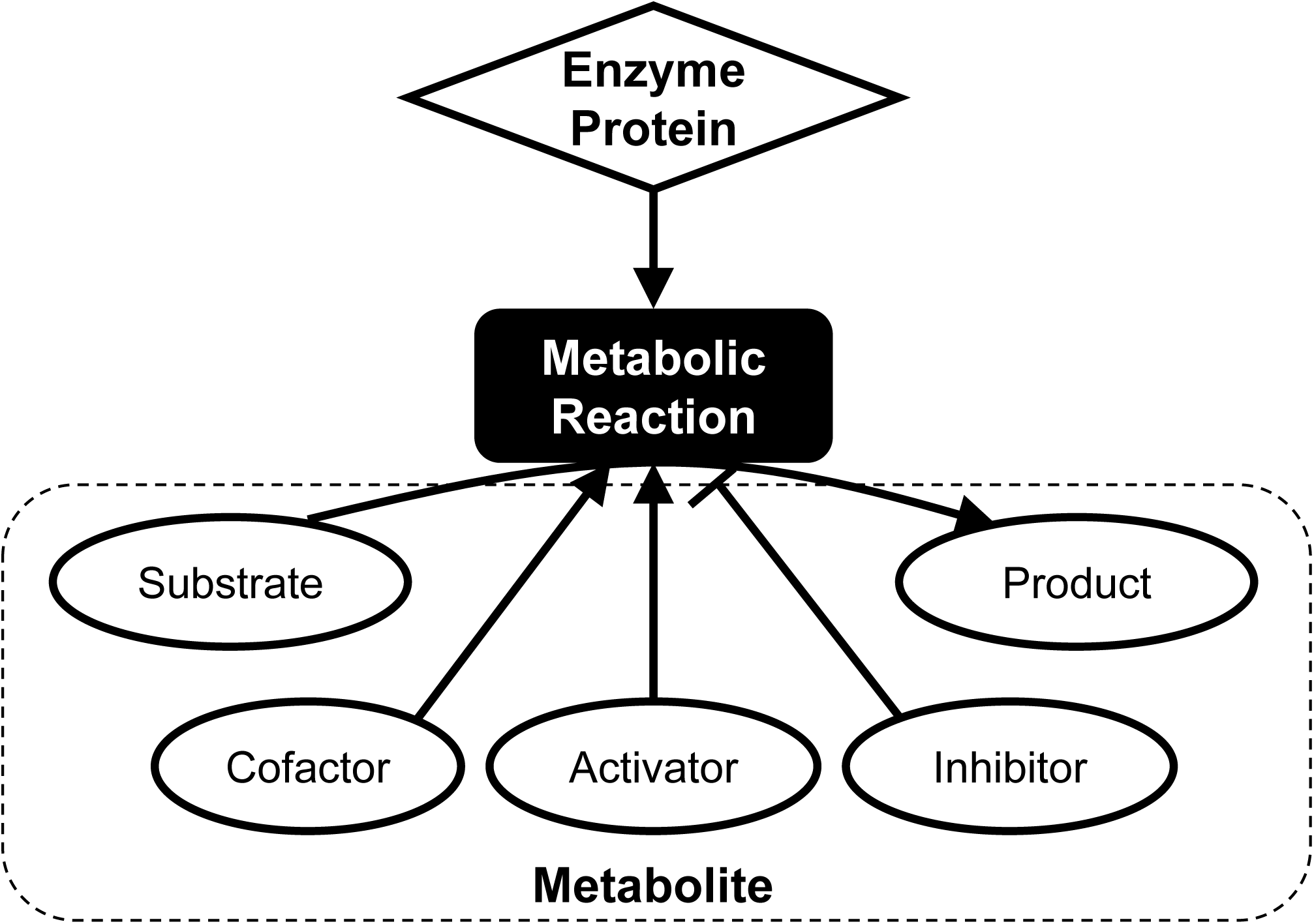

**fig. S3.**
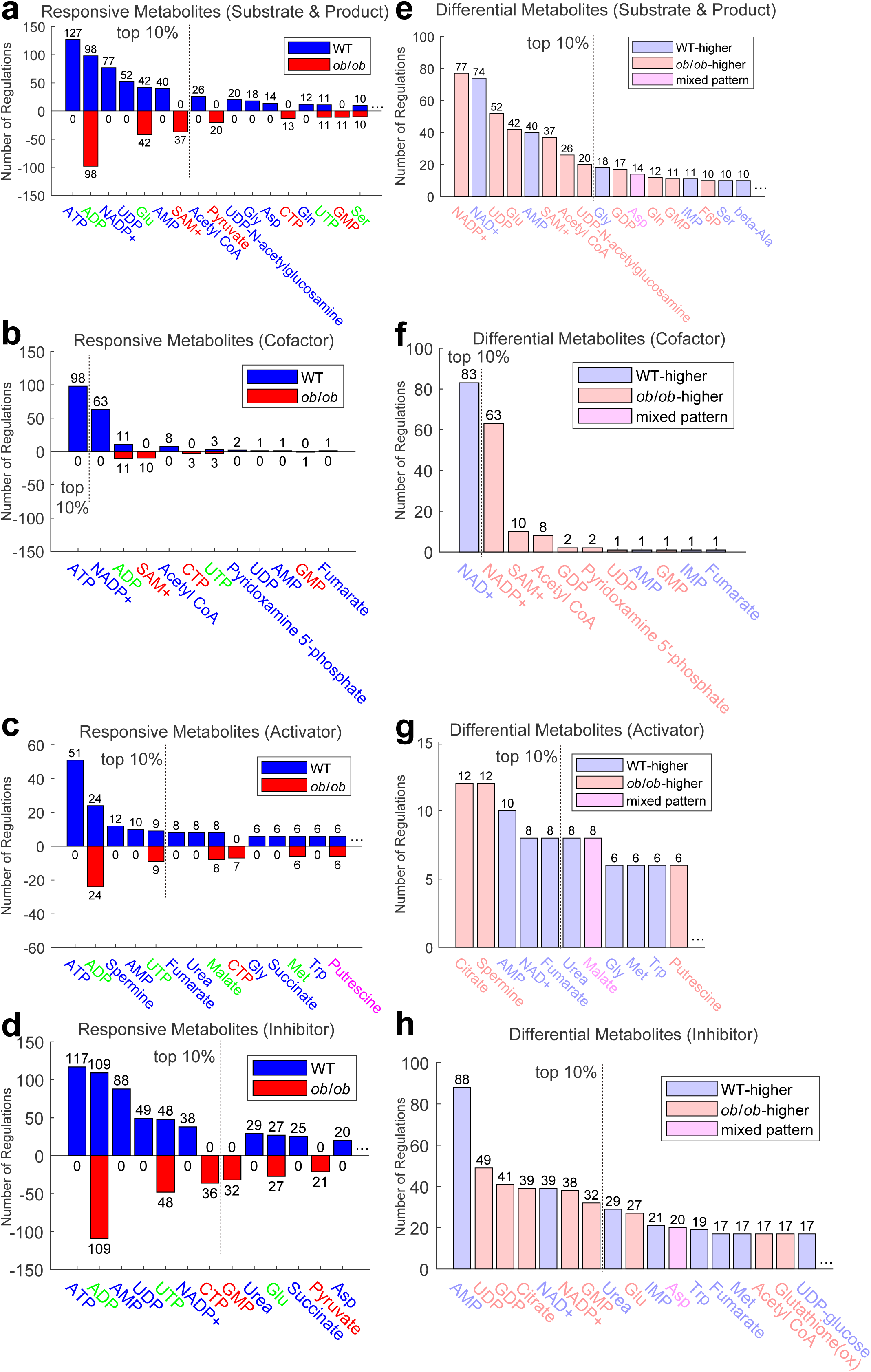

**fig. S4.**
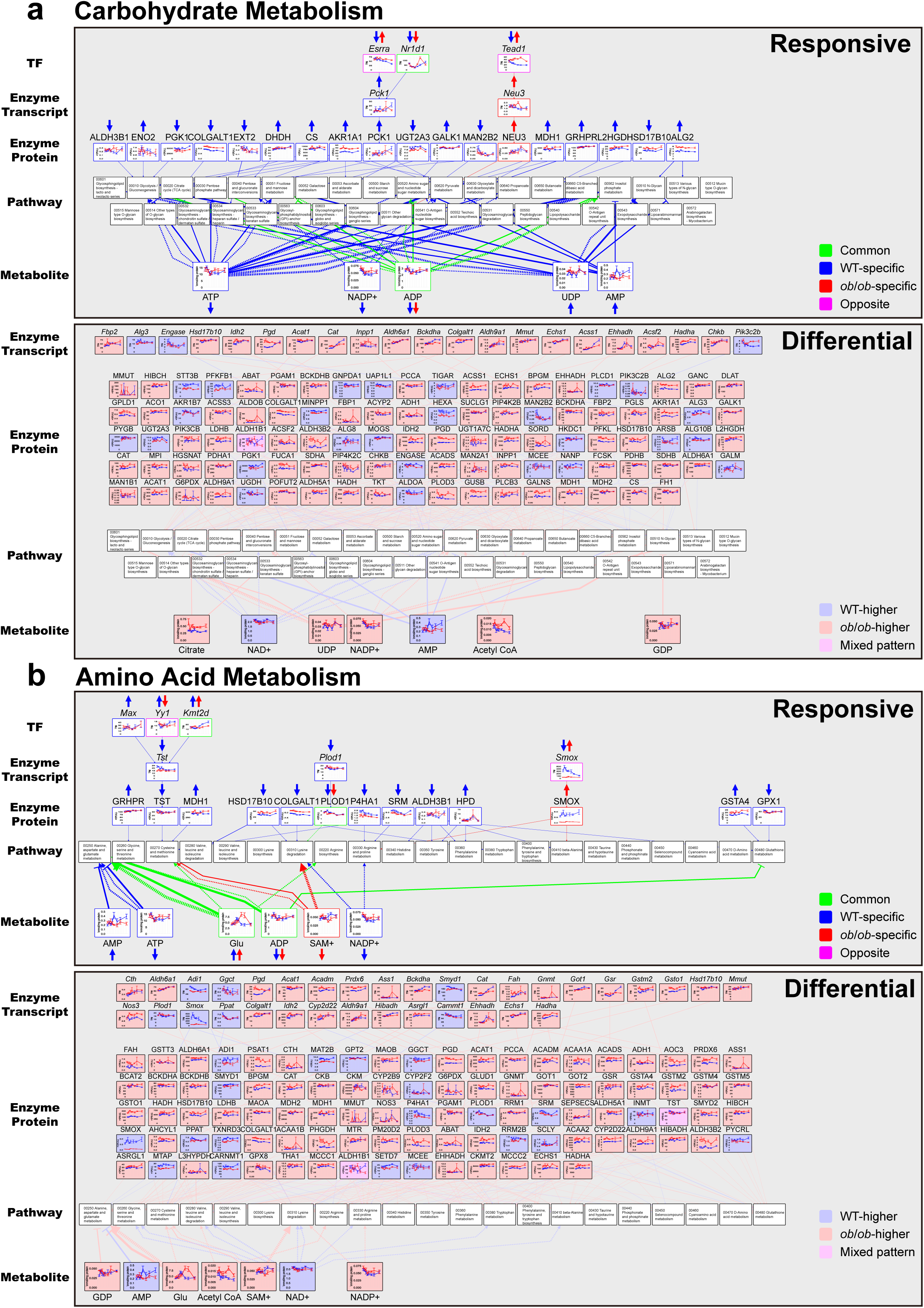
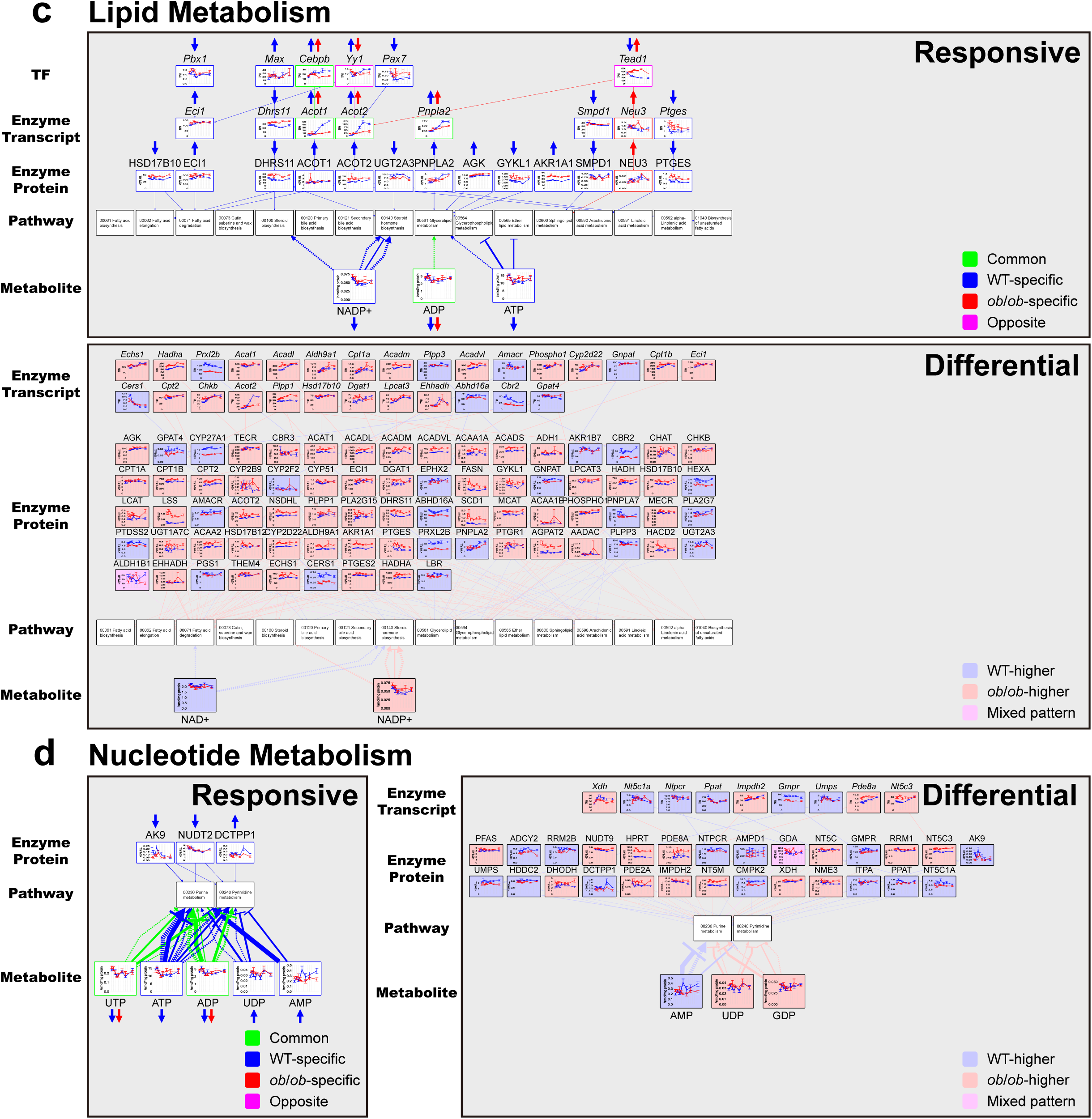

**fig. S5.**
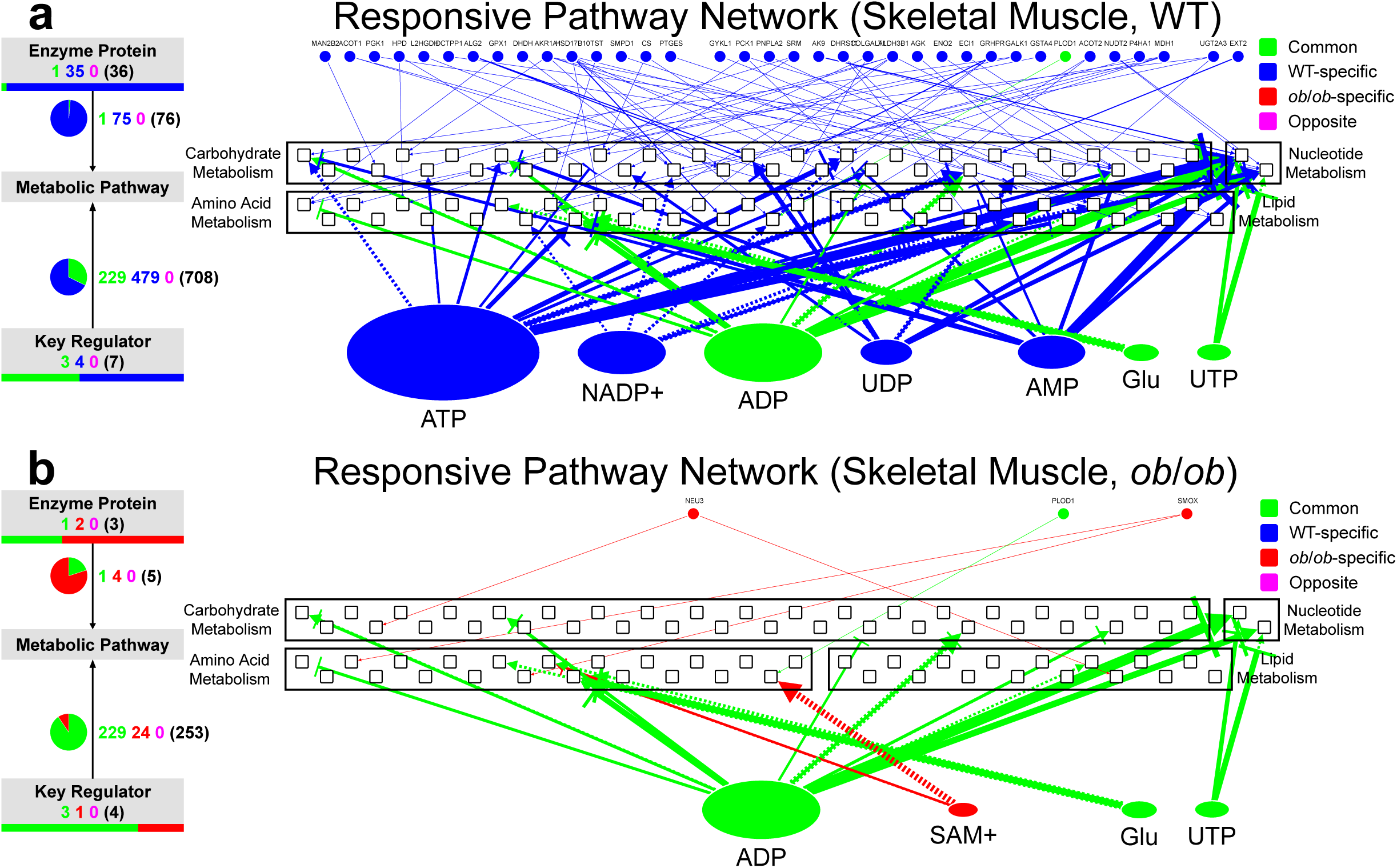

**fig. S6.**
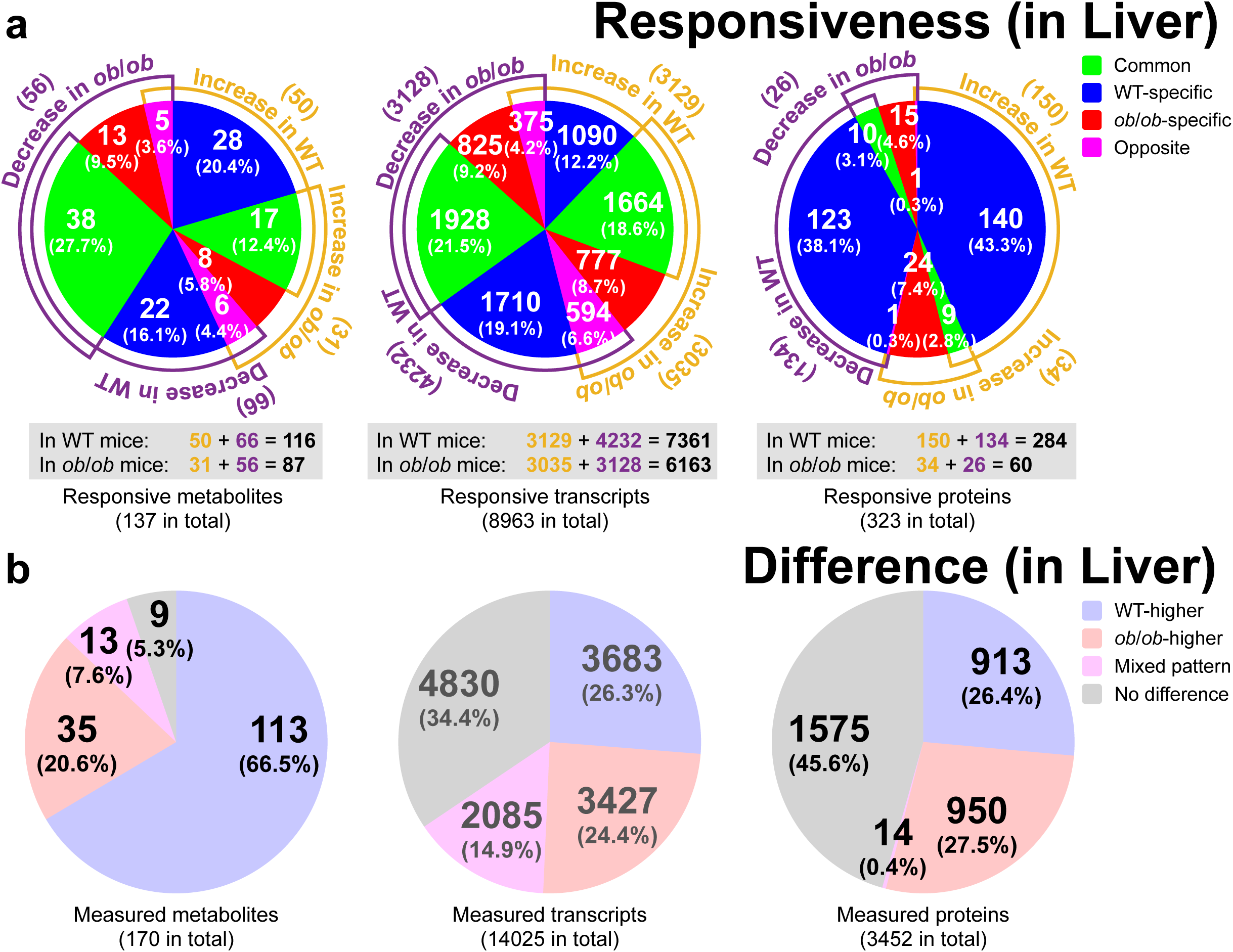

**fig. S7.**
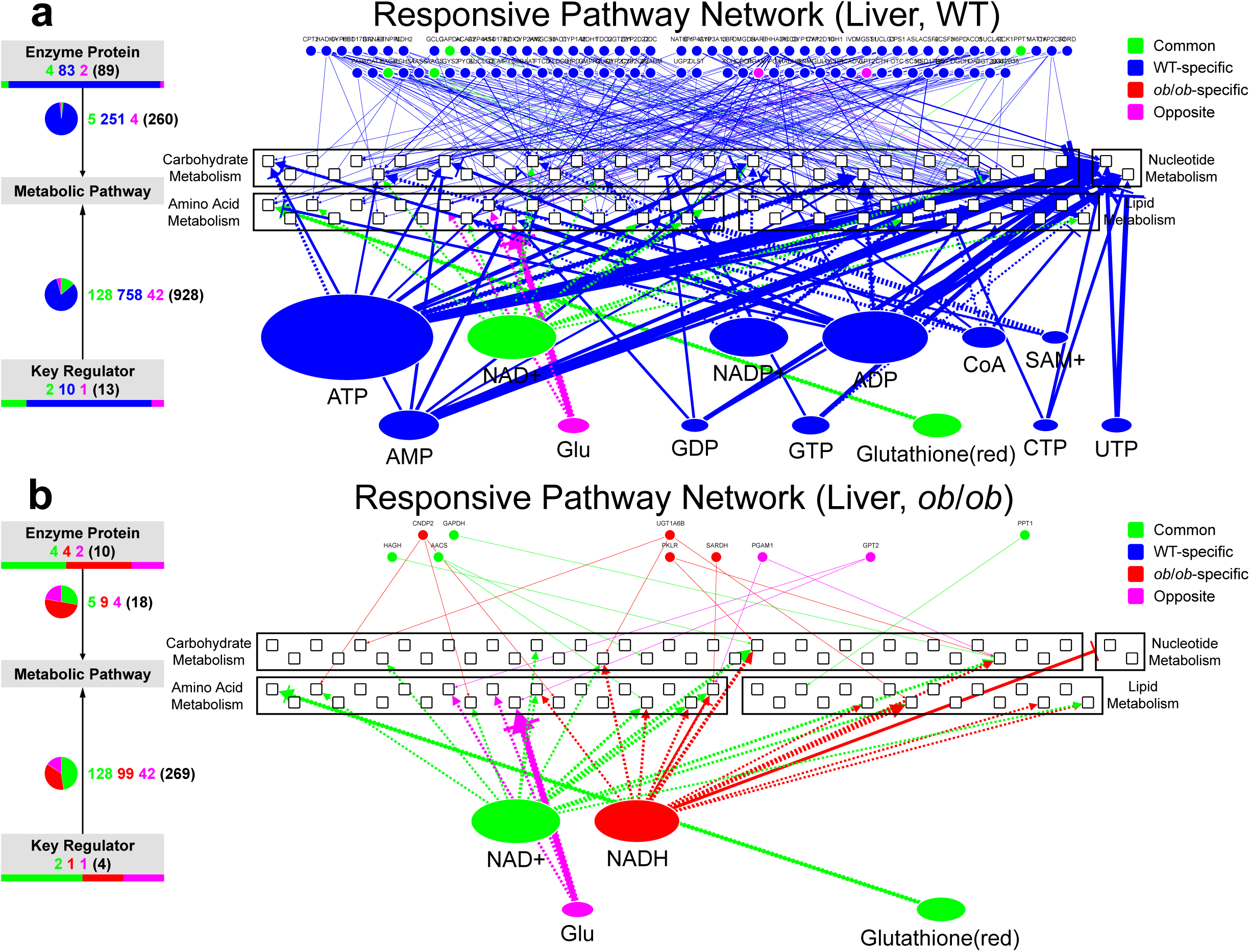

**fig. S8.**
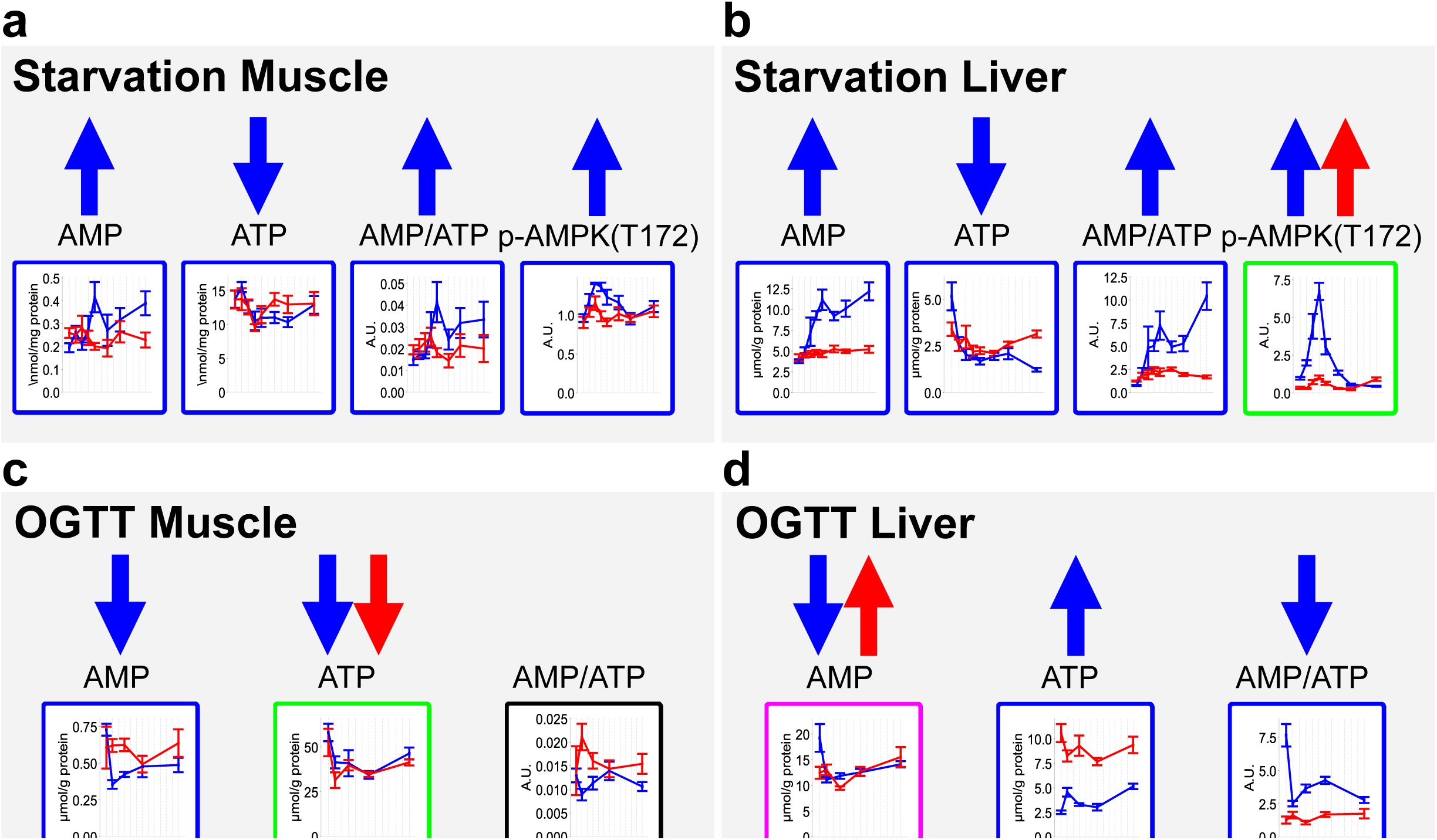

**fig. S9.**
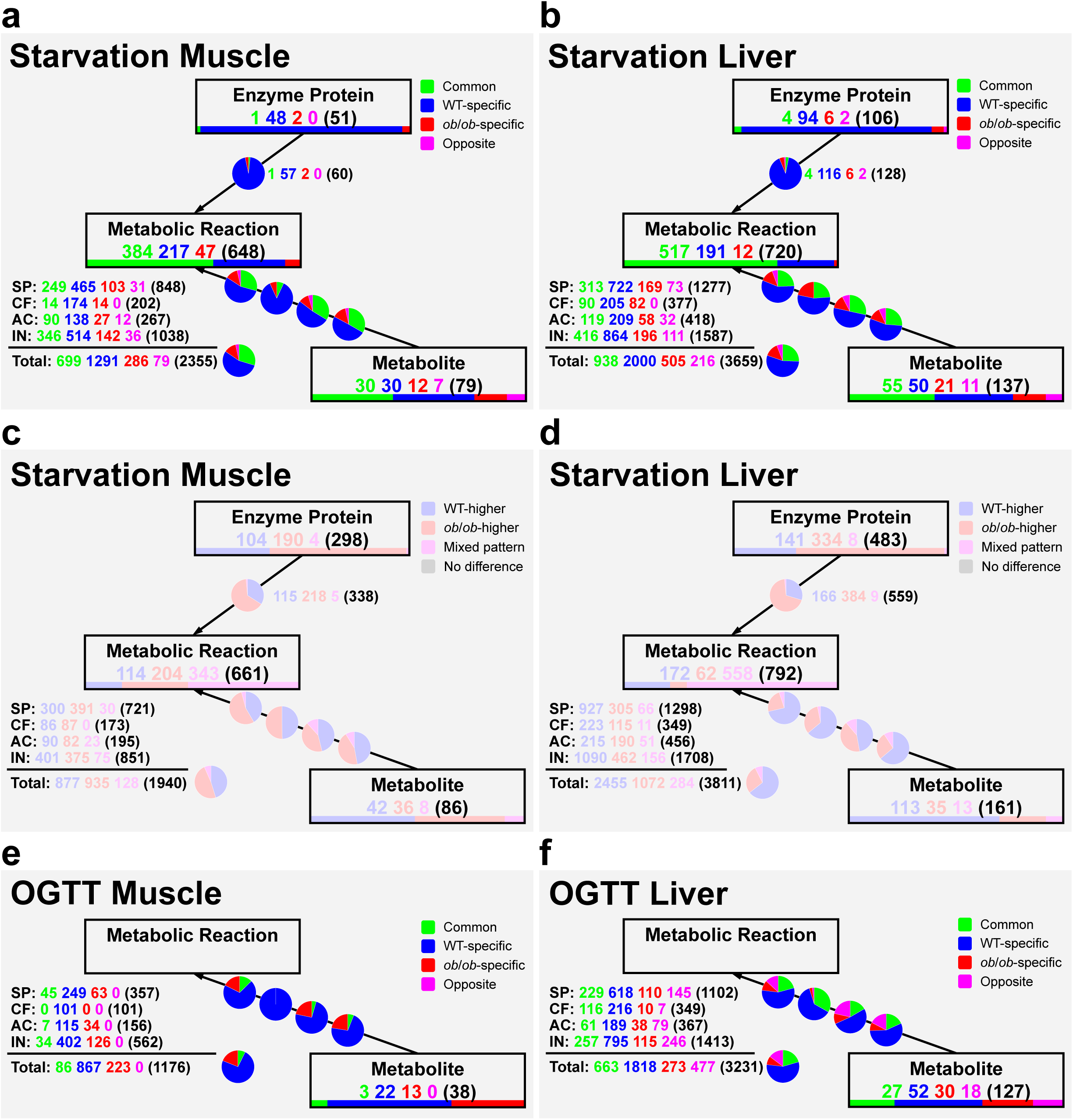

**fig. S10.**
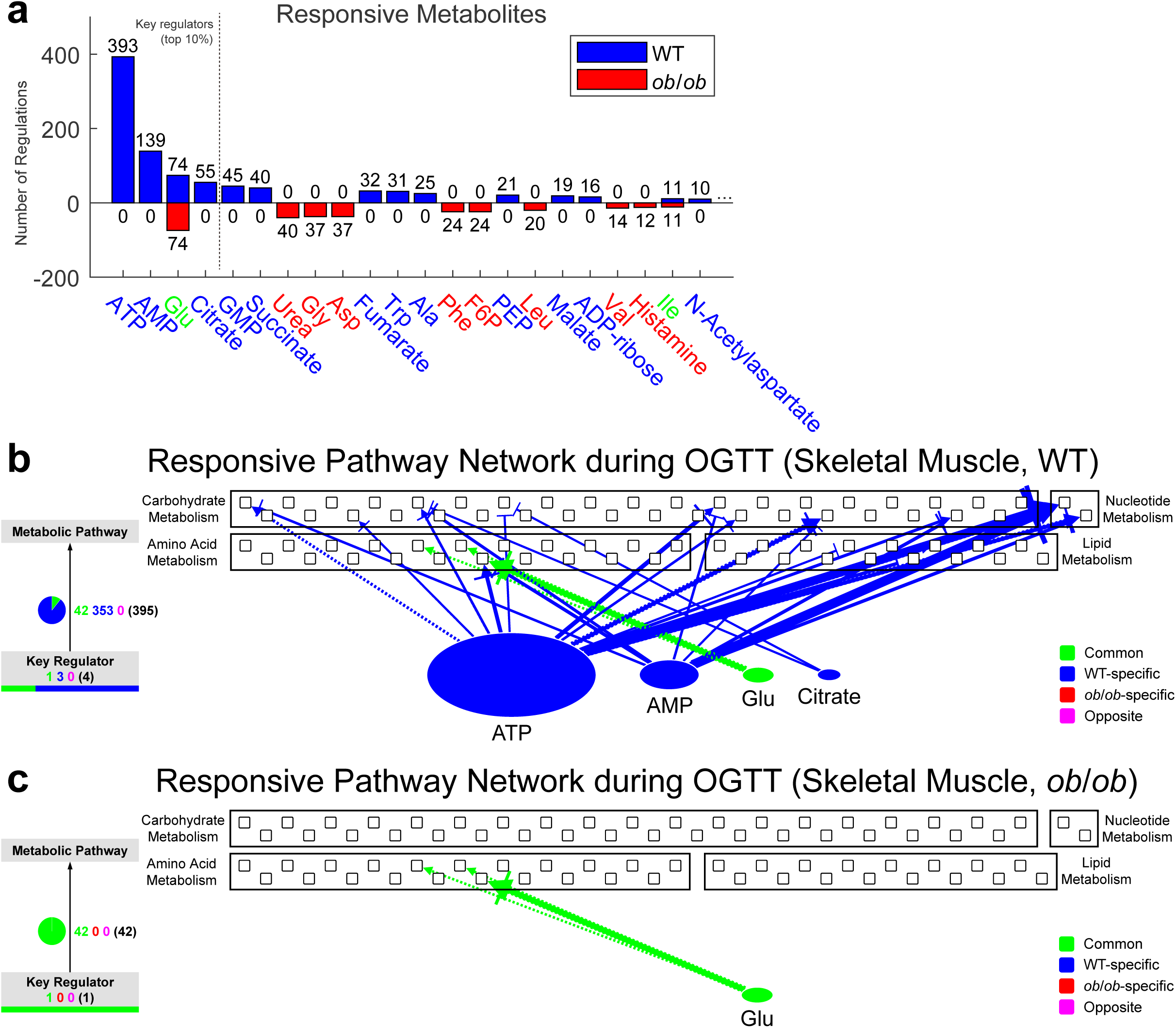

**fig. S11.**
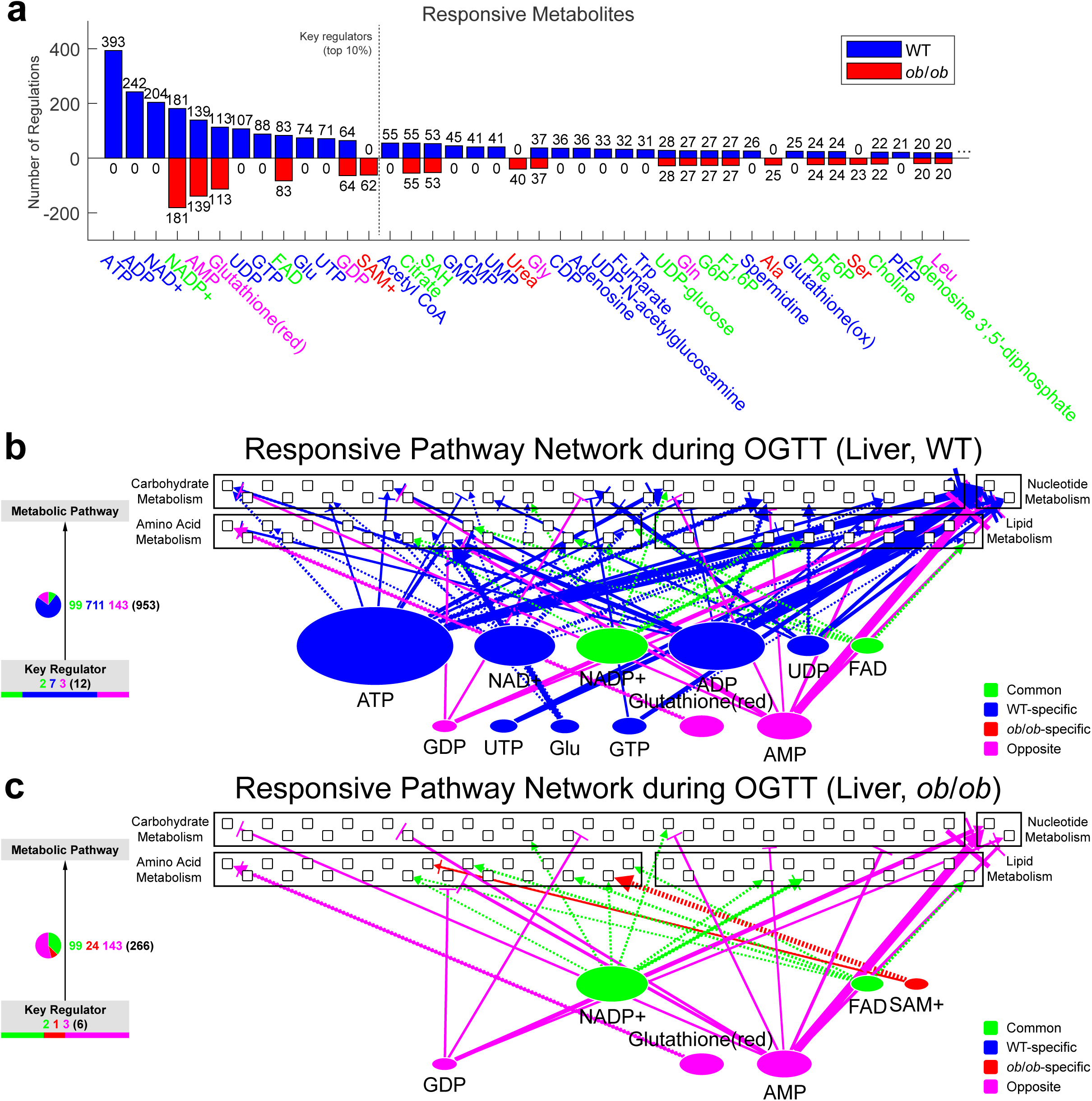

**fig. S12.**
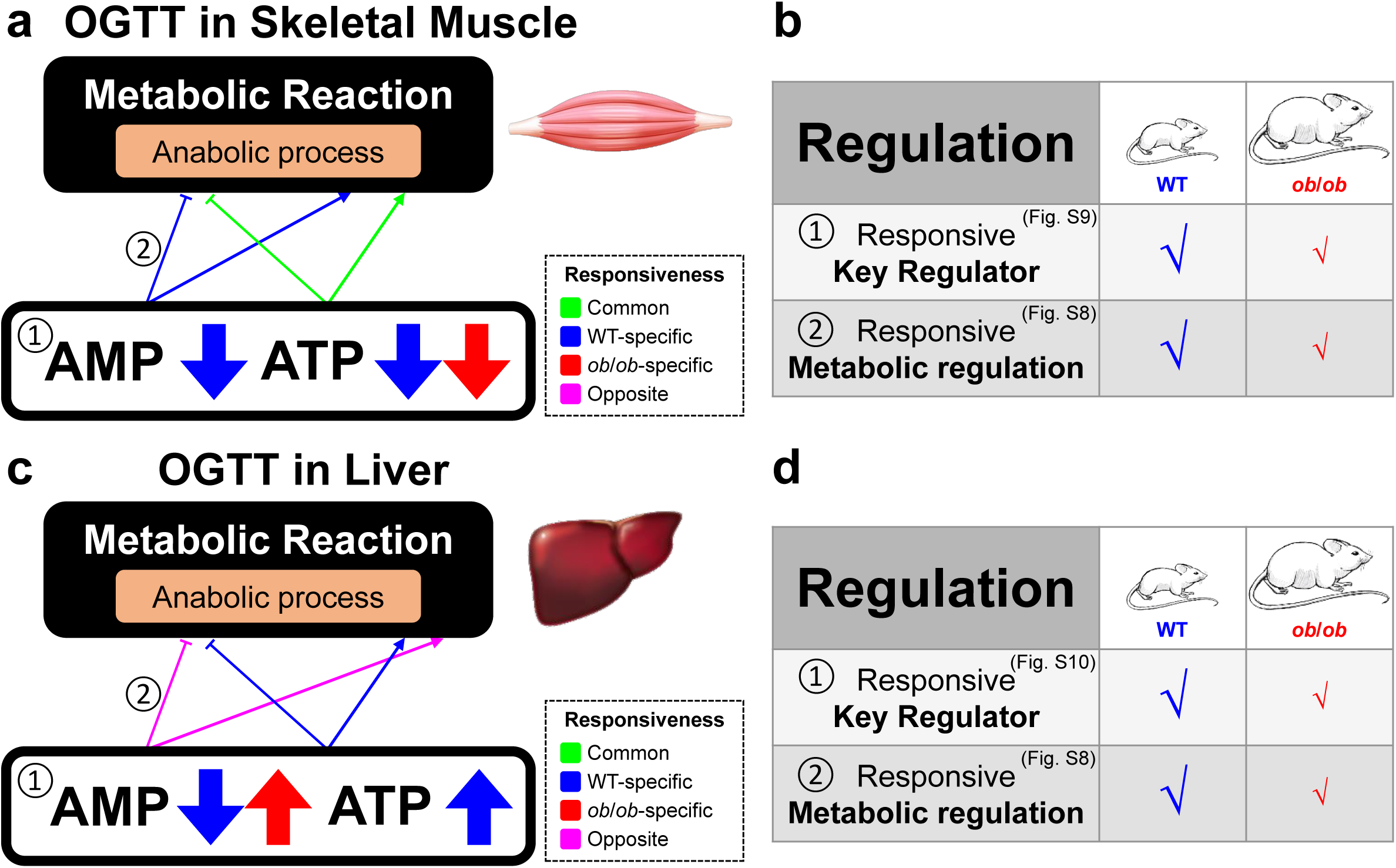

**fig. S13.**
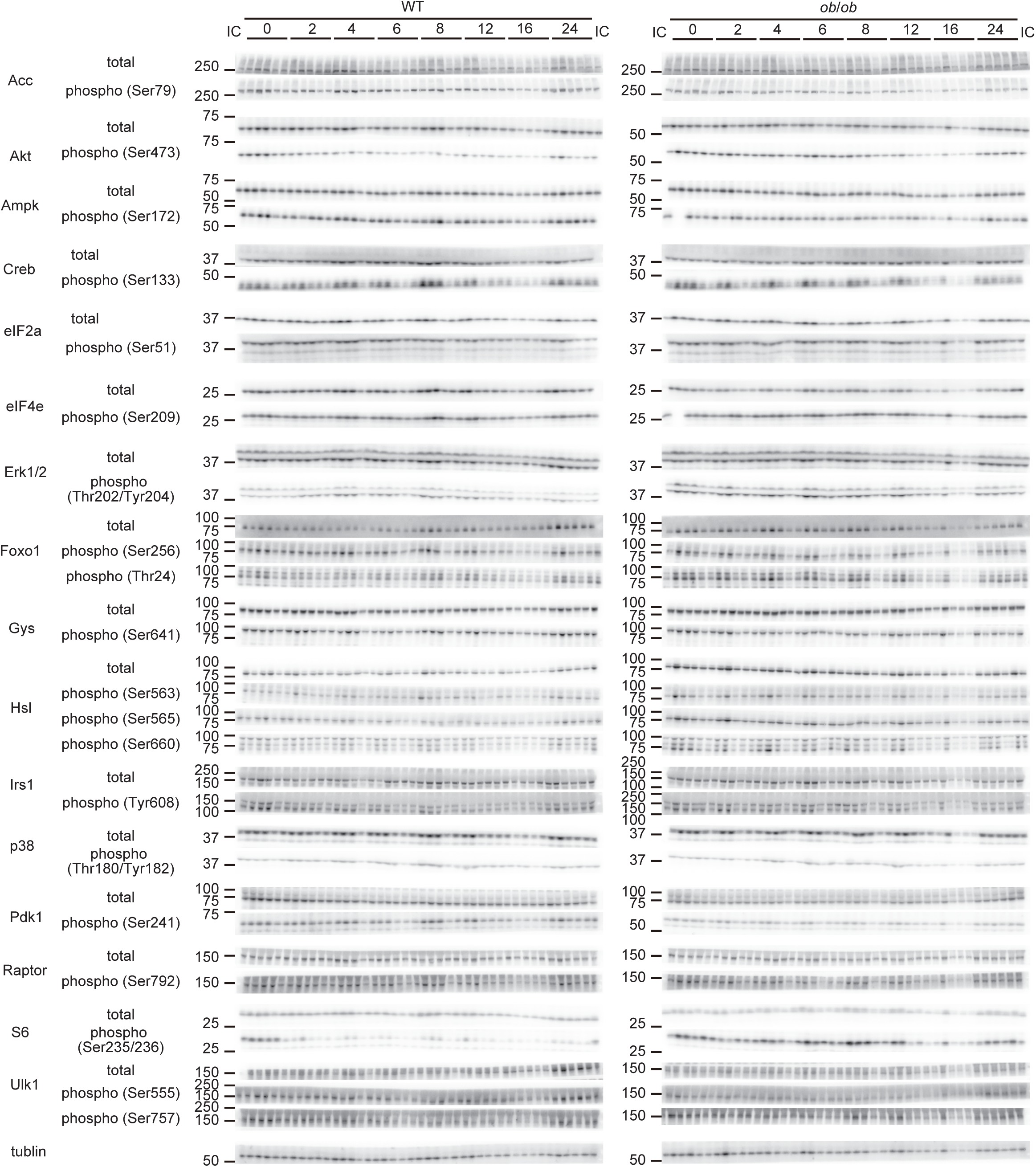

