## Supplementary material for "Global loss of responsiveness in key regulator metabolites and elevated enzyme proteins as metabolic dysregulation in skeletal muscle and liver of obese mice during starvation": Legends for Supplementary Figures

**SUPPLEMENTARY FIGURE TITLES AND LEGENDS**

**fig. S1: Blood insulin and blood glucose in WT and *ob*/*ob* mice during starvation**

(a-b) Blood insulin (a) and blood glucose (b) were measured at each time point during starvation. Time-course data are shown as means ± SD, with blue representing WT mice and red representing *ob*/*ob* mice. p < 0.05 was considered as responsive. *n* = 5 biological replicates per group.

**fig. S2: Nodes representing metabolic reactions in global networks and their regulation**

Metabolic reactions are represented as nodes, connected by their regulators, including enzyme proteins and metabolites. Metabolites that regulate metabolic reactions function as substrates, products, cofactors, allosteric activators, and allosteric inhibitors.

**fig. S3: Starvation-responsive and -differential key regulator metabolites with multifunctional roles**

(a-d) Numbers of regulations mediated by responsive metabolites functioning as substrates & products (a), cofactors (b), activators (c), and inhibitors (d). Blue bars above zero indicate regulations in WT mice, while red bars below zero indicate regulations in *ob*/*ob* mice. Metabolite name colors denote responsiveness: common (green), WT-specific (blue), *ob*/*ob*-specific (red), and opposite (magenta). Only the top responsive metabolites are shown in (a, c-d). (e-h) Numbers of regulations mediated by differential metabolites functioning as substrates & products (e), cofactors (f), activators (g), and inhibitors (h). Bar and metabolite name colors denote differences: WT-higher (light blue), *ob*/*ob*-higher (light red), and mixed pattern (light magenta). Only the top differential metabolites are shown in (e, g-h).

**fig. S4: Individual pathway networks for energy metabolism pathways**

(a-d) Starvation-responsive and -differential pathway networks for carbohydrate (a), amino acid (b), lipid (c), and nucleotide (d) metabolism. In responsive networks, node frame colors indicate responsiveness, while in differential networks, node fill colors indicate differences. Edge thickness represents the relative number of regulations mediated by each regulator. Time-course data are shown as means ± SEM, with blue representing WT mice and red representing *ob*/*ob* mice. In starvation-responsive networks, upward (increase) and downward (decrease) arrows denote responsiveness in WT (blue) and *ob*/*ob* (red) mice. *n* = 5 biological replicates per group.

**fig. S5: Individual pathway networks in skeletal muscle of WT and *ob*/*ob* mice during starvation**

(a-b) Starvation-responsive pathway network showing the regulation of carbohydrate, amino acid, lipid, and nucleotide metabolism by responsive enzyme proteins and key regulator metabolites in the skeletal muscle of WT (a) and *ob*/*ob* mice (b) during starvation. The numbers of responsive enzyme proteins and key regulator metabolites are represented by the horizontal stacked bar on the left. Node sizes and edge thickness reflect the number of regulations mediated by each responsive key regulator metabolite.

**fig. S6: Single-omic analysis of starvation-responsive and -differential metabolites, transcripts, and proteins in liver**

(a) Pie charts showing responsiveness patterns of metabolites, transcripts, and proteins. Numbers of increased (orange) and decreased (purple) molecules are indicated in brackets around the pie charts. Below the pie charts, the total numbers of responsive molecules in WT and *ob*/*ob* mice (sum of increased and decreased molecules) are shown, with the overall number of responsive molecules (WT and *ob*/*ob* combined) provided at the bottom. Responsiveness was defined as q < 0.1, determined using one-way ANOVA with BH correction. *n* = 5 biological replicates per group. (b) Pie charts showing difference patterns of metabolites, transcripts, and proteins. Total numbers of measured molecules are indicated in brackets at the bottom. Difference was defined as q < 0.1 at any time point, determined using one-way ANOVA with BH correction. *n* = 5 biological replicates per group.

**fig. S7: Individual pathway networks in the liver of WT and *ob*/*ob* mice during starvation**

(a-b) Starvation-responsive pathway network showing the regulation of carbohydrate, amino acid, lipid, and nucleotide metabolism by responsive enzyme proteins and key regulator metabolites in the liver of WT (a) and *ob*/*ob* mice (b) during starvation. The numbers of responsive enzyme proteins and key regulator metabolites are represented by the horizontal stacked bar on the left. Node sizes and edge thickness reflect the number of regulations mediated by each responsive key regulator metabolite.

**fig. S8: Comparison of the responsiveness of AMP, ATP, and AMP/ATP ratio in different conditions**

(a-d) Time course plots and responsiveness (indicated by arrows) of AMP, ATP, and the AMP/ATP ratio in skeletal muscle (a) and liver (b) during starvation, and in the skeletal muscle (c) and liver (d) during OGTT. Phosphorylation levels of AMPK are also shown for skeletal muscle (a) and liver (b) during starvation. Time-course data are presented as means ± SEM, with blue representing WT mice and red representing *ob*/*ob* mice. One-way ANOVA was used to compare across time points, and q-values were calculated using the BH correction. Responsiveness was defined as q < 0.1. *n* = 5 biological replicates per group.

**fig. S9: Comparison of global networks in the skeletal muscle and liver during starvation and OGTT**

(a-b) Calculation of regulation of metabolic reactions by responsive enzyme proteins and metabolites in the skeletal muscle (a) and liver (b) during starvation. (c-d) Calculation of regulation of metabolic reactions by differential enzyme proteins and metabolites in the skeletal muscle (c) and liver (d) during starvation. (e-f) Calculation of regulation of metabolic reactions by responsive metabolites in the skeletal muscle (e) and liver (f) during OGTT. In (a-f), node numbers are shown in boxes and represented by horizontal stacked bars below each box, while edge numbers are displayed as pie charts with corresponding labels. SP: Substrates/products; CF: cofactors; AC: allosteric activators; IN: allosteric inhibitors.

**fig. S10: Individual pathway networks in the skeletal muscle of WT and *ob*/*ob* mice during OGTT**

(a) Numbers of regulations mediated by responsive metabolites in the skeletal muscle during OGTT. Blue bars above zero indicate regulations in WT mice, while red bars below zero indicate regulations in *ob*/*ob* mice. Metabolite name colors denote responsiveness: common (green), WT-specific (blue), *ob*/*ob*-specific (red), and opposite (magenta). Only the top responsive metabolites are shown. (b-c) Starvation-responsive pathway network showing the regulation of carbohydrate, amino acid, lipid, and nucleotide metabolism by responsive key regulator metabolites in the skeletal muscle of WT (b) and *ob*/*ob* mice (c) during OGTT. The numbers of responsive key regulator metabolites are represented by the horizontal stacked bar on the left. Node sizes and edge thickness reflect the number of regulations mediated by each responsive key regulator metabolite.

**fig. S11: Individual pathway networks in the liver of WT and *ob*/*ob* mice during OGTT**

(a) Numbers of regulations mediated by responsive metabolites in the liver during OGTT. Blue bars above zero indicate regulations in WT mice, while red bars below zero indicate regulations in *ob*/*ob* mice. Metabolite name colors denote responsiveness: common (green), WT-specific (blue), *ob*/*ob*-specific (red), and opposite (magenta). Only the top responsive metabolites are shown. (b-c) Starvation-responsive pathway network showing the regulation of carbohydrate, amino acid, lipid, and nucleotide metabolism by responsive key regulator metabolites in the liver of WT (b) and *ob*/*ob* mice (c) during OGTT. The numbers of responsive key regulator metabolites are represented by the horizontal stacked bar on the left. Node sizes and edge thickness reflect the number of regulations mediated by each responsive key regulator metabolite.

**fig. S12: Summary of the distinct regulatory patterns in WT and *ob*/*ob* mice during OGTT**

(a-b) Summary of key regulatory strategies in the skeletal muscle of WT and *ob*/*ob* mice during OGTT. (a) Simplified network illustrating the regulation of metabolic reactions by responsive key regulator metabolites. (b) Table summarizing the primary differences between WT and *ob*/*ob* mice, with smaller check marks indicating that the regulation is present but less pronounced. (c-d) Summary of key regulatory strategies in the liver of WT and *ob*/*ob* mice during OGTT. (c) Simplified network illustrating the regulation of metabolic reactions by responsive key regulator metabolites. (d) Table summarizing the primary differences between WT and *ob*/*ob* mice, with smaller check marks indicating that the regulation is present but less pronounced.

**fig. S13: Western blot images**

Western blot results include measurements of both phosphoprotein and total protein for each phosphorylation event. Each run contained 40 samples from either WT or *ob*/*ob* mice, with two internal control (IC) lanes on each side. *n* = 5 biological replicates per group.
